# Evolution of iGluR ligand specificity, polyamine regulation, and ion selectivity inferred from a placozoan Epsilon receptor

**DOI:** 10.1101/2024.06.25.600656

**Authors:** Anhadvir Singh, Boris S. Zhorov, Luis A. Yanez-Guerra, Alessandra Aleotti, C. Defne Yanartas, Yunqi Song, Adriano Senatore

## Abstract

Epsilon ionotropic glutamate receptors (iGluRs) belong to a recently described sub-family of metazoan receptors that is distinct from the AMPA, Kainate, Delta, and Phi (*i.e.*, AKDF) sub-family, the NMDA sub-family, and the Lambda subfamily. Here, we sought to better understand the evolutionary and functional properties of Epsilon receptors by focusing on homologues from the basal invertebrate *Trichoplax adhaerens* (phylum Placozoa). We provide an updated species-guided phylogeny of eukaryotic iGluRs, and a comprehensive phylogeny of placozoan receptors uncovering marked diversification of Epsilon receptors within three conserved subclades, and four invariable subclades of AKDF receptors. Detailed functional characterization of the *T. adhaerens* Epsilon receptor GluE1αA revealed robust activation by glycine, alanine, serine, and valine, but not glutamate. Through combined of structural modeling and mutation experiments, we used GluE1αA to test the hypothesis that only a small set of amino acids in the ligand binding domain determine ligand selectivity. Mutation of just three amino acids converted GluE1αA selectivity to glutamate, resulted in nascent sensitivity to AMPA, and increased sensitivity to the AMPA/Kainate receptor blocker CNQX. Lastly, combined modeling and mutation experiments revealed that an atypical serine residue in the pore NQR site of GluE1αA, along with an aspartate four amino acids downstream, confers sensitivity to voltage-dependent polyamine block, while the serine alone diminishes both polyamine block and Ca^2+^ permeation compared to asparagine and glutamine residues of AMPA and Kainate receptors. Altogether, we demonstrate conserved molecular determinants for polyamine regulation between Epsilon and AKDF receptors, and evidence that natural variations in NQR residues have important implications for ion permeation and regulation by polyamines.

## Introduction

Ionotropic glutamate receptors (iGluRs) belong to a large and diverse family of ligand-gated ion channels present in eukaryotes and prokaryotes [1, 2]. The most studied of these are the vertebrate NMDA, AMPA and Kainate receptors, collectively known for their important roles in excitatory synaptic signaling in the mammalian brain [3, 4]. Recently, a landmark phylogenetic study uncovered two novel types of metazoan iGluRs, the Epsilon receptors that originated very early during animal evolution and are found broadly in bilaterians and non-bilaterians, and Lambda receptors that are exclusive to sponges (phylum Porifera) [5]. This study also revealed that AMPA and Kainate receptors belong to a larger group that also includes Delta and Phi receptors (hence collectively named AKDF receptors), and that NMDA receptors are absent in non-bilaterian animals except cnidarians. These findings have been corroborated by recent phylogenetic analysis, which have also reported that Lambda receptors are more phylogenetically proximal to plant receptors rather than other metazoan iGluRs [1, 6, 7]. To date, the functional properties of Lambda receptors have not been reported, but Epsilon receptors from the ctenophore species *Mnemiopsis leidyi* [8] and the basal chordate *Branchiostoma lanceolatum* [5] have been cloned and expressed *in vitro*, revealing unique properties including variable sensitivity to the amino acid transmitters glutamate and glycine. An interesting hypothesis that emerged from this and other comparative work, leveraging functional and structural studies observations of vertebrate and non-vertebrate iGluRs with bound ligands (reviewed in [3, 9]), is that only a small set of amino acids within the ligand binding domain (LBD) determine ligand selectivity [8, 10]. The most prominent ligands in discussion are glutamate and glycine, where predictions of ligand selectivity based on these noted amino acids point to frequent evolutionary switches in ligand selectivity within all major classes of metazoan iGluRs [10]. Indeed, although not yet experimentally validated, the hypothesis can explain why switches in ligand selectivity, particularly between glutamate and glycine, occurred so frequently during iGluR evolution [9, 10]. Another notable feature of the cloned Epsilon receptors is that their macroscopic currents undergo rectification at depolarized voltages, a hallmark feature of AMPA and Kainate receptors attributable to voltage-dependent block by endogenous cytoplasmic polyamines [11]. Nonetheless, whether Epsilon receptors are indeed modulated by polyamines, in a homologous manner as AKDF receptors, has not yet been confirmed.

An interesting group of animals in the context of metazoan iGluR evolution are the early-diverging placozoans, which possess AKDF and Epsilon receptors but lack NMDA receptors present only in cnidarians and bilaterians. These animals are millimetre-sized marine invertebrates that lack synapses and a nervous system but can coordinate their cells for directed locomotive behaviors including positive and negative chemotaxis [12–15], gravitaxis [16], and thermotaxis [17]. They also elicit behavioral responses to applied substances including glycine, glutamate, GABA, protons, and synthetic regulatory peptides that mimic endogenously encoded peptides [14, 15, 18, 19]. Presumably, these soluble ligands are detected on cell surfaces leading to downstream cytoplasmic responses, for example contraction or altered ciliary beating which drives placozoan locomotion. Accordingly, in addition to AKDF and Epsilon receptors, placozoans express a rich set of genes associated with neural signaling and detection of extracellular stimuli, including voltage-gated sodium, calcium, and potassium channels, mechanically gated channels such as TRP and Piezo, G protein coupled receptors, and other ligand-gated channels including P2X and Degenerin/Epithelial sodium channels [20–24]. To date, several studies have leveraged the basal phylogenetic position of placozoans to explore the functional properties and evolution of such genes, including voltage-gated calcium (Ca_V_) channels [25–27], Degenerin/Epithlial sodium channels [28, 29], and one AKDF receptor [30], all from the species *Trichoplax adhaerens*. Interestingly, the latter AKDF receptor was found to be highly atypical in conducting constitutive leak currents that are blocked by a broad range of amino acids including glycine, alanine, and D-serine, attributed to a unique tyrosine residue in the ligand binding domain.

In this study, we sought to better explore the evolutionary properties of placozoan iGluRs, first by generating a comprehensive phylogenetic analysis of iGluRs. This analysis, which includes sequences from four placozoan species (*T. adhaerens*, *Trichoplax* species H2, *Hoilungia hongkongnesis*, and *Cladtertia collaboinventa*), provides some advances in our general understanding of iGluR phylogeny in eukaryotes and uncovers previously undescribed clades of placozoan Epsilon receptors, promoting us to propose a new naming scheme based on these identified clades. We also conducted a detailed functional characterization of a *T. adhaerens* Epsilon receptor, named GluE1αA, bearing a non-canonical serine residue in the pore NQR site. Through a combination of electrophysiology, pharmacology, mutation analysis, and structural modeling, we provide evidence that GluE1αA is negatively regulated by polyamines, requiring an acidic aspartate residue four amino acids downstream of the NQR site. However, the NQR serine residue diminishes the extent of polyamine block while also reducing Ca^2+^ permeation, while mutation to canonical residues of glutamine, asparagine, and arginine increase polyamine block and Ca^2+^ permeation. Through a combination of mutation analysis and structural modeling, we also confirm the hypothesis that just a small set of defined amino acids in the ligand binding domain determine ligand selectivity, converting GluE1αA from a glycine activated, glutamate insensitive receptor, to one that is exclusively gated by glutamate. Furthermore, these same mutations altered the pharmacological properties of the receptor, establishing a nascent sensitivity to AMPA, and significantly increasing sensitivity to the AMPA/Kainate receptor blocker CNQX.

## Results

### Eukaryotic phylogeny of iGluRs and a revised nomenclature of placozoan iGluRs

We performed phylogenetic analysis to explore the evolutionary relationships of placozoan iGluRs within a broader phylogenetic context of metazoans and non-metazoan eukaryotes (Figure 1a and Supplementary Figures 1 to 4). To overcome the challenges related to inferring phylogenetic relationships between sequences of a single gene family from a wide range of distantly related species, we performed gene tree to species tree reconciliation (see Materials and Methods). Such species-tree-aware inference leverages species relationships to better resolve uncertain branch positions in a gene tree, and discriminates between duplication (D) and speciation (S) events at each branch [31]. Moreover, reconciling the gene tree with the species tree allows to root the resulting tree based on the information of species relationships. According to our analysis, animal-specific iGluRs include a broad AKDF group, an Epsilon group which is paralogous to the AKDF receptors, and a NMDA group, which is paralogous to both AKDF and Epsilon receptors (Figure 1a and Supplementary Figure 1). Ancient duplication events appear to have given rise to AMPA, Kainate, and Phi receptors in a bilaterian ancestor of protostomes and deuterostomes. While AMPA and Kainate receptors are well represented in both deuterostomes and protostome invertebrates, Phi receptors are less common in protostomes with a single clade of lophotrochozoan receptors and a single receptor from the ecdysozoan/arthropod species *Calanus glacialis* (Supplementary Figure 1). For comparison, a previous phylogenetic analysis did not identify Phi receptors outside of deuterostomes [5], while a more recent analysis similarly identified a clade of Phi receptors in Lophotrochozoans [1]. Our additional identification of a single Phi receptor from the arthropod *C. glacialis* thus supports deep ancestry of Phi receptors in protostome invertebrates but extensive losses in ecdysozoans (Supplementary Figure 2).

**Fig. 1:**
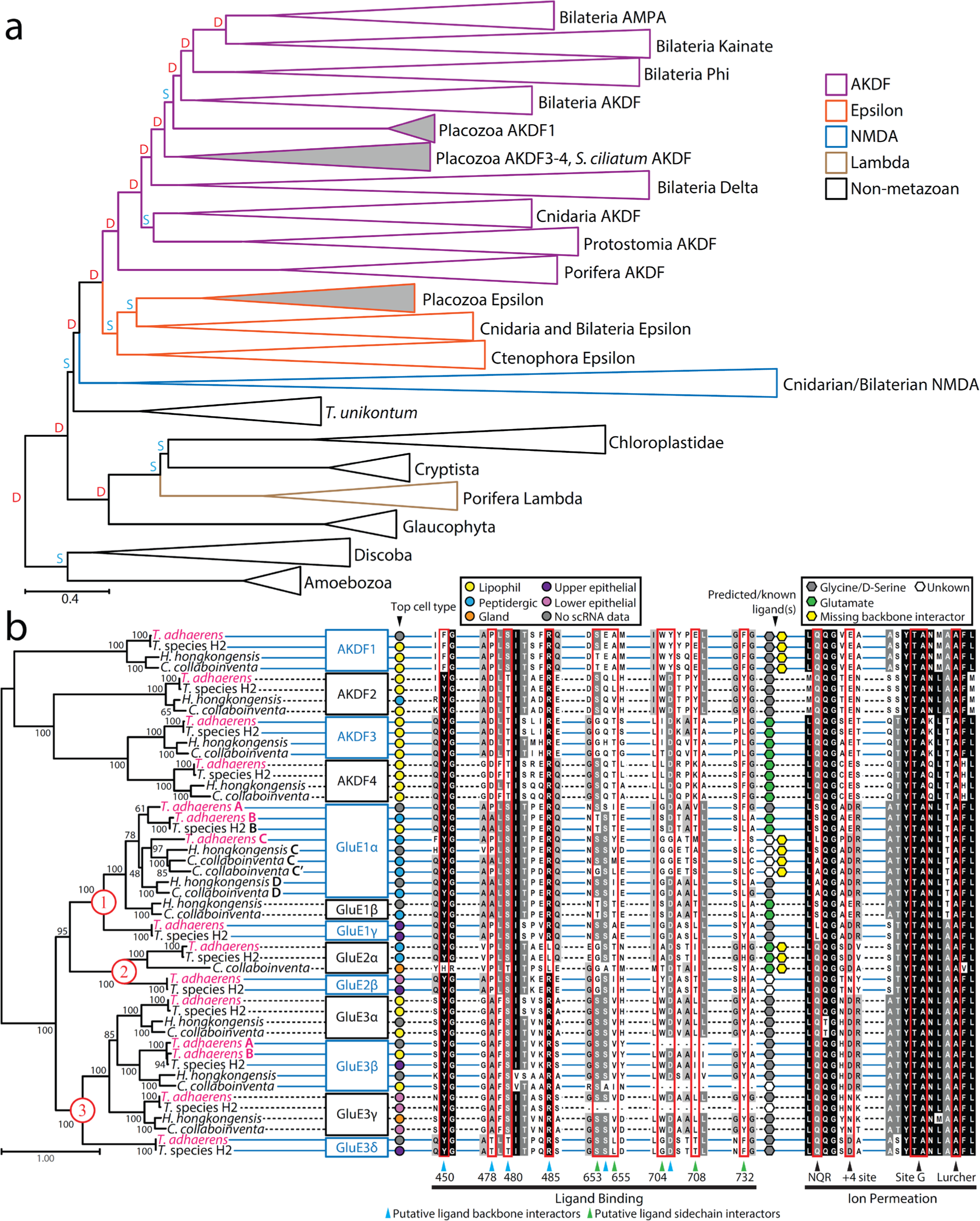
Phylogeny and revised nomenclature of placozoan iGluRs. **a** Species aware phylogenetic tree of identified iGluR proteins sequences from select eukaryotes and metazoans. The letters D (red) and S (cyan) denote predicted duplication and speciation and events, respectively, predicted by the GeneRax software [31]. **b** Maximum likelihood phylogenetic tree of placozoan iGluRs reveals invariable AKDF orthologues and a diversity of epsilon receptors falling within three major clades (1 to 3 labeled in red font). The black numbers on the nodes of the tree indicate ultrafast bootstrap values for 1000 replicates. Top cell type expression based from single cell RNA-Seq data [24], predicted ligand selectivity for glycine/D-serine vs. glutamate are indicated by symbols to the right of the tree according to the corresponding legends, and aligned amino acids sequences associated with ligand binding and ion permeation are indicated below the alignments. Horizontal solid blue and dashed black lines running behind the aligned residues are used to demark different clades of placozoan iGluRs according to our proposed nomenclature indicated on the right of the phylogenetic tree.

Our analysis also supports a common origin of Delta iGluRs in a basal bilaterian ancestor, finding phylogenetic homologues in deuterostome and lophtrochozoans from the phyla Mollusca and Annelida, consistent with previous studies [1, 5, 32]. Notably, one of these studies also identified a single Delta receptor from the cnidarian species *Nematostella vectensis* (GluAKDF1 with accession number v1g50912) [1], suggesting an even deeper phylogenetic origin. However, this same protein sequence (accession number XP_001633354.1 in our analysis) formed a strong and separate clade with an expanded set of cnidarian AKDF receptors in both our gene tree and our species-reconciled tree (Figure 1a, Supplementary Figures 1 and 3). Our analysis also identified two clades of placozoan AKDF receptors, both positioned between Delta receptors and a superclade of bilaterian AKDF receptors the separate into group I (containing AMPA, Kainate, and Phi receptors), and group II (comprised of uncharacterized receptors from molluscs, annelids, cephalochordates, and the hemichordate *Ptychodera flava*). The earliest branching clade of placozoan receptors includes a single receptor from the calcareous sponge *Sycon ciliatum*, consistent with a previous report [1].

With respect to Epsilon type receptors, we corroborate their existence in protostome invertebrates [1] by identifying numerous homologues from the marine annelid *Capitella teleta*, contrasting earlier studies that found them only in deuterostomes [5, 6]. Our analysis also supports, as previously suggested, a pre-bilaterian origin of type 1 NMDA receptors, present also in cnidarians, and a bilaterian origin of NMDA2 and NMDA3 receptors, emerging via duplication of an ancestral NMDA2/3-like receptor after the cnidarian/bilaterian divergence [1, 5, 6] (Supplementary Figure 1). Also consistent with previous studies, we find that the sponge-specific Lambda receptors cluster outside of metazoan iGluRs [1, 6, 7], associating with a set of distant iGluRs from several species within the Diaphoretickes (Supplementary Figures 1 and 2). This association is also apparent in a clustering analysis where the Lambda receptors associate with these non-metazoan receptors and not metazoan iGluRs (Supplementary Figure 3). Notably, while we identified Lambda receptors in the gene data for species within the sponge clades Calcarea and Homoscleromorpha, we did not do so in the data for the three demosponge species included in our analysis (Table S1). Two possible explanations for these various observations are that: 1) Lambda receptors underwent strong sequence divergence in Porifera causing them to incorrectly associate with non-metazoan receptors, followed by gene loss in select clades within Porifera, or 2) as previously suggested [1], Lambda receptors were passed from the Diaphoretickes lineage to a sub-clade of sponges via lateral gene transfer.

Although most clades in our species aware tree correspond with those in our iGluR gene tree, several clades differ between the two reflecting alterations incurred during species reconciliation (Supplementary Figure 4). One of these is a clade of placozoan AKDF type 1 receptors, that in the gene tree associates with a set of deuterostome receptors, while in the reconciled tree it forms a sister relationship with bilaterian group I and II AKDF receptors (Figure 1a and Supplementary Figure 4). Another alteration is apparent for a clade of protostome AKDF receptors, which have an orthologous relationship with a set of cnidarian AKDF receptors in the reconciled tree but are included with the bilaterian Delta receptors in the gene tree. A third reconciliation is apparent for a clade of receptors from the distant unicellular lineages of Amoebozoa and Discoba, which have a paralogous relationship with NMDA receptors in the gene tree but are sister to all other eukaryotic iGluRs in the reconciled tree (Figure 1a and Supplementary Figure 4). Lastly, there is a slight difference in the position of Lambda receptors from sponges (Porifera), although in both the species and gene tree, they fall outside of metazoan iGluRs instead associating with iGluRs from species within Diaphoretickes.

Next, we re-examined the phylogeny of placozoan iGluRs using a more focused, manual approach for sequence identification. Specifically, we used BLAST and manual annotation to comprehensively identify iGluR sequences in the gene data available for the four placozoan species *Trichoplax adhaerens* [20–22], *Trichoplax* species H2 [22], *Hoilungia hongkongensis* [23], and *Cladtertia collaboinventa* [24], and used these sequences to generate a maximum-likelihood protein phylogeny (see Materials and Methods). The resulting tree revealed one-to-one orthology of AKDF receptors 1 to 4, contrasting the Epsilon receptors with instances of lineage-specific duplication or loss producing more diversity between the four placozoan species (Figure 1b). The comprehensive nature of our phylogenetic analysis, which delineates three major clades of placozoan Epsilon receptors, prompted us to propose a new nomenclature scheme based on phylogenetic groupings. Specifically, while AKDF1 to 4 receptors retain their previous name, we propose naming the three clades of placozoan Epsilon receptors GluE1 to GluE3. *T. adhaerens* possesses four GluE1 paralogues, three within a strongly supported clade comprised of homologues from all other species (*i.e.*, GluE1α subtypes A to C), and a fourth, also found in *Trichoplax* species H2 (GluE1γ), forming a sister relationship with other GluE1 sequences. Several nodes within the GluE1 clade are poorly supported, making it difficult to infer how these receptor subtypes evolved. However, it is notable that *H. hongkongensis* and *C. collaboinventa*, which are more phylogenetically related to each other [24], uniquely share GluE1β receptor orthologues, while *T. adhaerens* and *Trichoplax* species H2, also more closely related, uniquely possess GluE1γ receptors. In the GluE2 clade, there is one-to-one orthology of GluE2α and GluE2β receptors between *T. adhaerens* and *Trichoplax* species H2, and a single GluE2α orthologue from *C. collaboinventa* but none identified for *H. hongkongnesis*. More conservation is apparent in the GluE3 clade, with one-to-one orthologous relationships for GluE3α, GluE3β, and GluE3γ receptors. The exceptions are GluE3β, which uniquely duplicated in *T. adhaerens* giving rise to GluE3βA and GluE3βB, and GluE3δ, which we only found in the gene data from *T. adhaerens* and *Trichoplax* species H2. This proposed nomenclature, which should withstand incorporation of new sequences should they become available, is summarized in Table S2 along with corresponding accession numbers and previous names of relevant receptors.

We sought to determine whether the phylogenetic relationships of placozoan iGluRs are also reflected in by similar mRNA expression at the cellular level. Specifically, we mined the available single cell RNA-Seq data [24] to determine the top cellular expression of each receptor, revealing that the orthologous AKDF1 to AKDF4 receptors are enriched in digestive lipophil cells (Figure 1b), except for AKDF2 from *H. hongkongensis*, which is most abundant in neuron-like peptidergic cells despite also having strong expression in lipophil cells. In contrast, most of the GluE1 Epsilon receptors show enriched expression in peptidergic cells, with a notable exception being GluE1γ from *T. adhaerens* and *Trichoplax* species H2 which are enriched in upper/dorsal epithelial cells. More diversity is apparent within the GluE2 and GluE3 clades, though GluE3α and GluE3β show a general enriched expression in lipophil cells, and GluE3γ in lower/ventral epithelial cells.

We also predicted the ligand specificities of the various placozoan iGluRs based on aligned sequences that form key contacts with ligand amino acids in other receptors and are proposed to be deterministic for ligand preference (reviewed in [8, 10]). Specifically, while residues at positions 450, 478, 480, 485, 654, and 705 form interactions with ligand amino acid backbone atoms (numbered according to alignment with the mature rat GluA2 subunit with UniProt accession number P19491; Supplementary File 1), residues 653, 655, 704, 708, and 732 form contacts with side chain atoms and hence play key roles in defining selectivity. Thus, the presence of glycine, serine, or threonine at position 653, in conjunction with threonine at position 655 and tyrosine at position 732 is associated with glutamate specificity, while serine at position 653, along with a hydrophobic residue at 655 (valine, leucine, alanine, isoleucine, or proline) and phenylalanine at position 732, is associated with glycine/D-serine specificity [8, 10]. Based on these residues, placozoan AKDF1 and 2 receptors are predicted to bind glycine, while AKDF 3 and 4 are predicted to bind glutamate (Figure 1b). Notably however, the AKDF1 receptors lack an aspartate 750 (D_705_) residue, which is engaged in electrostatic interactions with the ligand alpha amino group and is important for ligand binding. Recently, this natural variation in sequence was shown to cause the *T. adhaerens* AKDF1 receptor to conduct constitutive leak currents that are blocked by a broad range of ligands, most pronounced for glycine [30]. A subset of GluE1α epsilon receptors are similarly missing the D_705_ residue, and have variable sequences at other key sites that prevents prediction of their ligand specificities (Figure 1b). The GluE2α orthologues also lack key backbone interacting residues, including arginine 485 (R_485_) which forms key electrostatic contacts with the ligand alpha carboxyl group but are predicted to bind glutamate. The consequences of these missing backbone-interacting residues in GluE1α and GluE2α receptors are unknown. Of the remaining epsilon receptors, extensive ligand switching is suggested in the GluE1 clade, where for example the duplicated paralogue GluE1αA from *T. adhaerens* is predicted to bind glycine, while GluE1αB is predicted to bind glutamate. Instead, most GluE3 receptors are predicted to bind glycine and D-serine, with a few receptors having divergent sequences that preclude prediction.

We also examined sequences in the pore-loop (P-loop) region located between the transmembrane helices M1 and M3 of each subunit, as well as the M3 helices which together with the P-loop govern ion permeation and selectivity. Most placozoan receptors possess a glutamine in the NQR site, a key locus that defines monovalent and divalent cation selectivity [3, 33], thus resembling AMPA and Kainate receptors (Figure 1b). GluE1 subunits are the exception having non-canonical residues of serine, alanine, or leucine at this position. All placozoan iGluR subunits, except GluE2β and GluE3γ, possess negatively charged aspartate or glutamate residues four amino acids down stream of the NQR site, which together with NQR residues contribute to polyamine binding in the pore and voltage-dependent block of ion permeation (reviewed in [11]). Lastly, considerable divergence is apparent in the M3 helix bearing the SYTANLAAF motif, which in the tetrameric quaternary structure of iGluRs forms a gate that moves outwards during activation [34]. Nonetheless, complete conservation is evident for alanine residues in the lurcher position (Figure 1b), which when mutated produce constitutively active channels [35], and alanine-threonine residues in site G, recently shown to be involved in Ca^2+^ permeation [36].

### The Trichoplax GluE1αA receptor has a broad ligand selectivity and moderately fast recovery from desensitization

Since the available transcripts for the *T. adhaerens* iGluRs were algorithmically assembled from RNA-Seq reads, we sought to verify their coding sequences by amplifying them in full-length (*i.e.*, start to stop codon) from a whole animal cDNA library via PCR, followed by cloning into the mammalian expression vector pIRES2-EGFP and sequencing in triplicate. This led to the successful cloning and verification of 10 iGluRs, whose sequences we submitted to GenBank (see Materials and Methods). When cloning the various iGluRs, we included a consensus mammalian Kozak sequence of GCCGCCACC upstream of the start codon allowing efficient expression of the channels in transfected Chinese Hamster Ovary (CHO)-K1 cells for whole-cell patch-clamp electrophysiology and functional characterization. In preliminary experiments, we were able to observe ligand-gated current for several AKDF and Epsilon receptors, including robust currents for the GluE1αA receptor expressed on its own and hence functional as a homotetrameric channel. For this first study, we decided to conduct a comprehensive functional analysis of GluE1αA. As predicted from sequences in the ligand-binding region (Figure 1a), GluE1αA is activated by glycine producing robust currents in CHO-K1 cells, also sensitive to D-serine, L-serine, and alanine, and weak activation by valine (Figure 2a). We observed no responses to glutamate and aspartate (Figure 2a), nor arginine, threonine, methionine, and betaine tested on their own at 10 mM, nor histidine, lysine, isoleucine, leucine, and phenylalanine tested as a mix each at 3 mM. Dose response curve analysis of activating ligands revealed strongest sensitivity to alanine, followed by glycine, L-serine, D-serine, all with EC_50_ values in the sub-millimolar range (*i.e.*, alanine = 0.049 ±0.011 mM, L-serine = 0.134 ±0.019 mM, and D-serine = 0.191 ±0.044 mM), followed by valine with an EC_50_ of 1.51±0.24 mM (Figure 2b and c, Supplementary Figure 5). Altogether, GluE1αA most resembles the vertebrate NMDA receptor subunit GluN1 and the ctenophore Epsilon subunit ML032222a in its ligand selectivity, as both also bind glycine, alanine, and serine [3, 8]. Instead, the Epsilon subunit from the basal chordate *Branchiostoma lanceolatum*, dubbed GluE1, is not sensitive to alanine, serine, or glutamate, despite being activated by glycine [5].

**Fig. 2:**
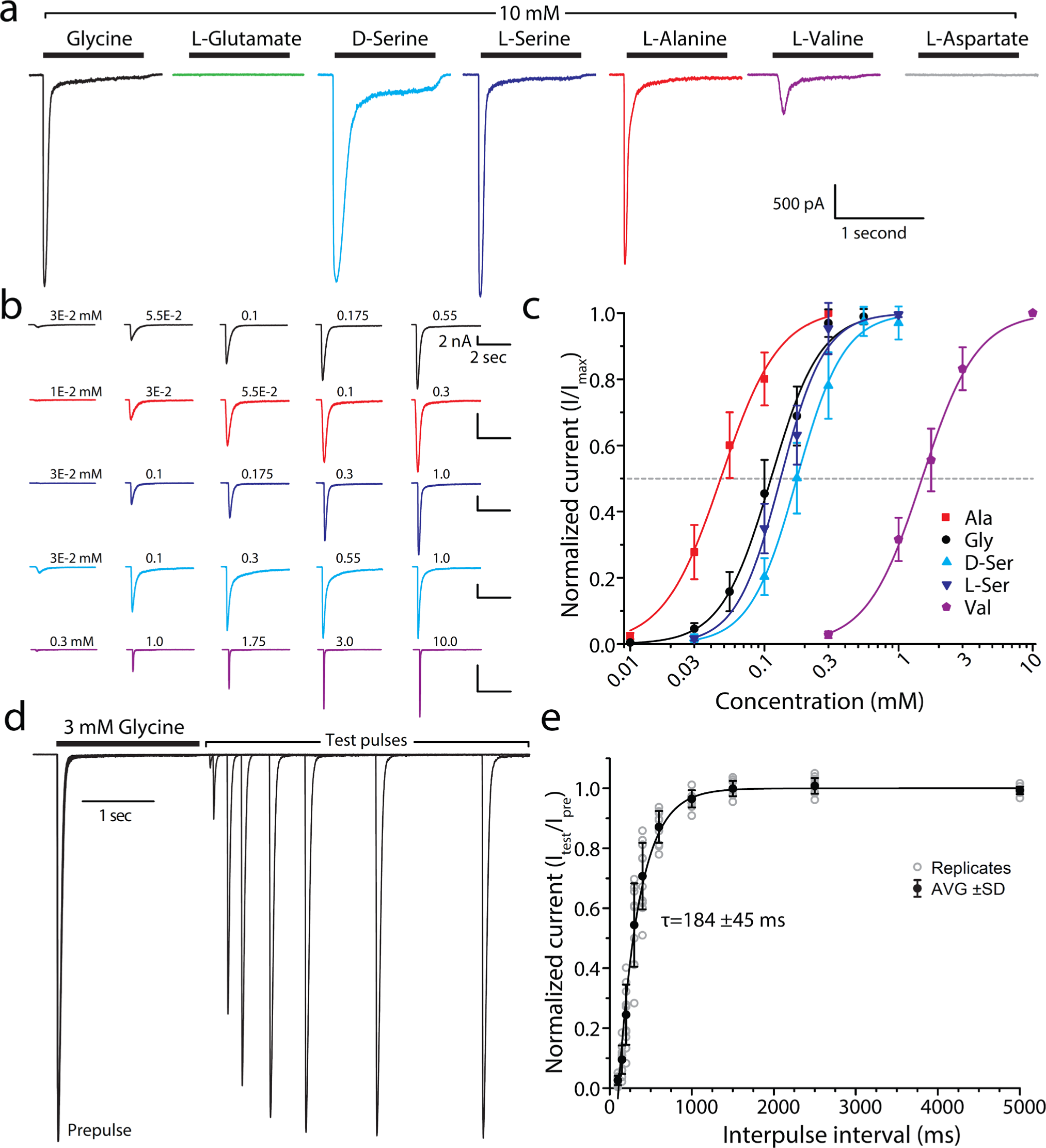
*In vitro* currents of the *T. adhaerens* GluE1αA epsilon receptor. **a** Sample whole-cell currents of the *T. adhaerens* GluE1αA receptor expressed in CHO-K1 cells upon extracellular perfusion of 10 mM glycine, glutamate, L-serine, D-serine, alanine, valine, and aspartate. **b** Sample recordings of GluE1αA receptor macroscopic currents elicited by increasing concentrations of amino acid ligands (top to bottom: glycine in *black*, alanine in *red*, L-serine in *navy blue*, D-serine in *light blue*, and valine in *purple*). **c** Dose response curves showing greatest sensitivity to alanine, and lowest sensitivity to valine. The EC_50_ values for all ligands were statistically different from glycine as shown by two-sample T-tests (p<0.05). **d** Sample paired pulse currents of GluE1αA elicited by 3 mM glycine applied for 2 seconds (to desensitize the channels), followed by a wash step of increasing duration then 2 second test pulse. **e** Plot of average recovery from desensitization of GluE1αA after prolonged application of 3 mM glycine, fitted to a single exponential with a τ value of 184 ± 45 ms (n=10).

Perfusion of a 2 second pulse of 3 mM glycine, followed by washes of increasing duration followed by a second 2 second pulse of 3 mM glycine, revealed fast monophasic recovery from desensitization for GluE1αA with a time constant of 184 ±45 milliseconds (Figure 2d and e). The *T. adhaerens* receptor is thus much faster in its recovery compared to the Epsilon receptor from the lancelet *Branchiostoma lanceolatum* with a time constant of 10.8 seconds [5], and from the ctenophore *Mnemiopsis leidyi* with a time constant of 81 seconds [8].

### Mutation of key residues in the ligand binding domain switches ligand selectivity from glycine to glutamate

As noted, a model has been put forward defining iGluR ligand selectivity based on a set of key amino acids in the ligand binding domain [8, 10]. This prediction proved to be true for the GluE1αA receptor (Figure 1b), which as predicted is insensitive to glutamate but activated by glycine and D-/L-serine (Figure 3a to c). To the best of our knowledge, whether these few positions indeed define ligand selectivity, and moreover, can be altered to switch specificity to natural ligands, has not been experimentally tested, although sequences in the LBD suggest such switches have occurred naturally and frequently during evolution, evident for example among both the AKDF and epsilon receptors of placozoans (Figure 1b). Thus, according to the ligand-selectivity model, we switched the serine and isoleucine residues at equivalent positions 653 and 655 of GluE1αA to glycine and threonine, respectively, to convert its ligand specificity to glutamate (Figure 3a). A third mutation, phenylalanine to tyrosine at position 732, was generated to make the *T. adhaerens* epsilon receptor like NMDA2A and AMPA2 from human, reasoning that these are selective for glutamate and hence this mutation might enhance glutamate sensitivity.

**Fig. 3:**
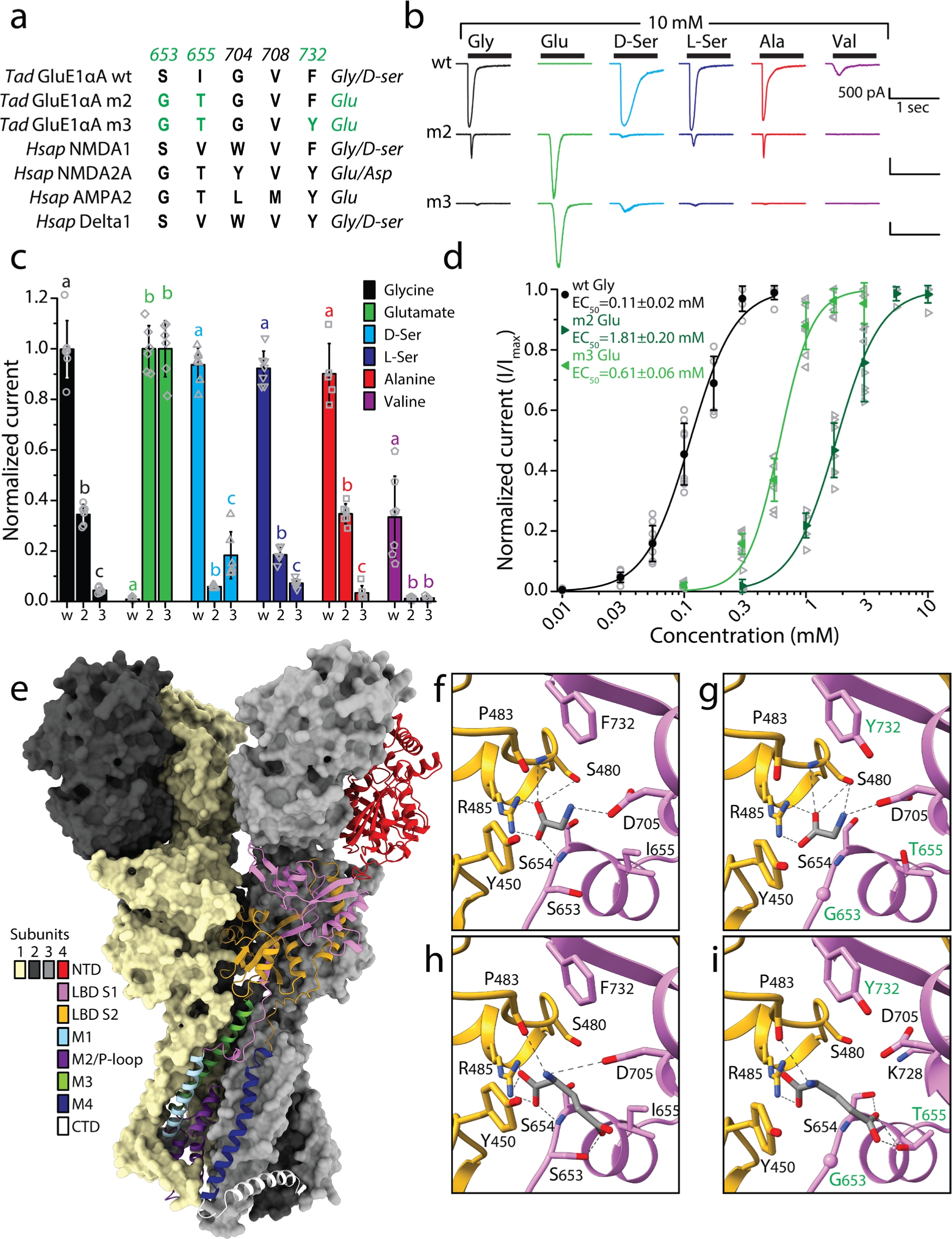
Mutation of key residues in the ligand binding domain of GluE1αA narrows its ligand-selectivity by switching to glutamate. **a** Multiple sequence alignment of deterministic residues that interact with ligand amino acid side chains and mediate ligand specificity in mammalian NMDA, AMPA, and Kainate receptors (numbers correspond to amino acids positions in rat GluA2). The residues in red were mutated in the wildtype (wt) GluE1αA receptor to resemble the glutamate-selective NMDA-2A and AMPA-2 receptors from human (the double and triple mutant variants are referred to as m2 and m3, respectively). **b** Sample whole-cell currents of wt, m2, and m3 variants of GluE1αA elicited by application of different amino acids at 10 mM. **c** Plot of average normalized peak inward currents of wt, m2, and m3 variants of GluE1αA in response to different amino acid ligands at 10 mM (n=6-7). Letters above the bars denote statistically significant differences among ligand types for post hoc Tukey tests (p<0.05) after One Way ANOVAs (glycine: F=295.00, p=2.26E-13; glutamate: F=284.71, p=1.18E-12; D-serine: F=679.87, p=3.33E-16; L-serine: F=209.44, p=3.7E-11; alanine: F=337.26, p=3.8E-14; valine: F=23.33, p=1.81E-5). **d** Dose response curves of wt GluE1αA normalized current responses to glycine (n=8), and of the m2 and m3 variants of to glutamate (n=6). One Way ANOVA confirmed significant differences among the EC_50_ values (F=350.89, p<1E-14), and Tukey post hoc tests revealed that all three were different from each other (p<5E-7). **e** AlphaFold 3-predicted quaternary structure of the GluE1αA tetramer. Three of the four subunits are shown as surfaces colored pale goldenrod, black, and gray, while the fourth subunit is depicted as a ribbon structure with different structural regions colored differently as indicated by the legend. NTD: N-terminal domain; LBD S1: ligand binding domain segment 1; LBD S2: ligand binding domain segment 2; M1 to M4: transmembrane helices 1 to 4; P-loop: pore-loop; CTD: C-terminal domain. **f** Homology modeling and docking of glycine in the putative ligand binding pocket of the wildtype GluE1αA receptor. **g** Homology modeling and docking of glycine in triple mutant (m3) variant of GluE1αA. **h** Homology modeling and docking of glutamate in the wt GluE1αA receptor. **i** Homology modeling and docking of glutamate in triple mutant (m3) variant of GluE1αA. In panels f to i, predicted hydrogen bonds are depicted by dashed grey lines.

Indeed, application of various activating ligands at 10 mM concentrations revealed that the double mutant variant of GluE1αA (dubbed m2, bearing the S_653_G and I_655_T mutations), became sensitive to glutamate with relatively diminished sensitivity to glycine, L-serine, D-serine, alanine, and valine (Figure 3b and c). The triple mutant (m3), bearing the additional F_732_Y mutation, almost completely lost sensitivity to these various amino acids in lieu of robust glutamate-activated currents. Dose response analysis revealed weaker affinity to glutamate for the m2 variant compared to glycine for the wildtype receptor (*i.e.*, glycine EC_50_ for the wildtype receptor = 0.11 ±0.02 mM, glutamate EC_50_ for the m2 variant = 1.81 ±0.20 mM), but inclusion of the F_732_Y mutation significantly increased glutamate sensitivity decreasing the EC_50_ to 0.61 ±0.06 mM (Figure 3d).

To gain insights into why these amino acid changes resulted in a ligand selectivity switch, we predicted the structure of GluE1αA with AlphaFold [37, 38] and conducted docking experiments of glycine and glutamate bound to the ligand binding pocket of the wildtype and m3 receptor variants. Interestingly, the AlphaFold 3 predicted tetrameric structure of the *T. adhaerens* epsilon receptor (Figure 3e) resembles the resolved structures of vertebrate homotetrameric GluD1 and GluD2 delta receptors, with the N-terminal domain (NTDs) and ligand binding domains (LBDs) forming a dimer of dimers arrangement, as opposed to the NTD-LBD swapped domains apparent for AMPA and Kainate receptors (reviewed in [4]). We then generated AlphaFold models of the isolated LBD and transmembrane domain (TMD) of wildtype and m3 variants of GluE1αA and docked glycine or glutamate with a starting position that mimicked the location of D-serine bound to the LBD of the receptor rat GluD2, determined via cryo-EM (PDB number 2v3u; [39]). Intensive Monte Carlo energy-minimizations yielded complexes shown in Figure 3f to I and Supplementary Figure 6. The energy scores of ligand-receptor interactions are provided in Table S3a to c. Glycine binding to the wildtype channel is stabilized mainly by salt bridges with residues D_705_ and R_485_ and non-bonded attraction to K_728_ and Y_450_ (Figure 3f; Supplementary Figure 6a; Table S3b). Of note, the total energy scores of glycine bound to the wildtype and m3 receptor variants reflect experimental observations, with the m3 variant having a higher energy score of -2.51 kcal/mol compared to -5.92 kcal/mol for wildtype (Table S3b), mainly due to weaker salt bridges and repulsive contributions from the mutated Y_732_ residue. Furthermore, Y_732_ is predicted to hydrogen bond with D_705_, shifting the latter away from the ligand and hence contributing to the lower affinity for glycine in the m3 LBD. The energy score of glutamate in the wildtype channel was positive (unfavorable) mainly due to repulsion from D_705_, S_480_, and E_656_, while a salt bridge between the glutamate ligand and R_485_ provides a strong attractive contribution. The interaction energy score of glutamate with the mutant channel was negative mainly due to attractive contributions from R_450_, K_728_, as well as the mutated residues G_653_, T_655_. Besides indicated residues, which provide major attractive or repulsive contributions to ligand-channel interactions, there are other residues that contribute energies with absolute value over 0.1 kcal/mol (Table S3c). For these analyses, it should be emphasized that the estimated interaction energy scores between ligands and the LBD were weak, indicating that these are sensitive to imprecision of the applied force field. Indeed, although our computations are consistent with the trend of ligand-channel energy changes, the predicted energy scores should not be considered as measures of ligand affinity.

### Mutations that alter ligand selectivity also affect sensitivity to vertebrate iGluR agonists and the AMPA/Kainate blocker CNQX

Since pharmacological agonists of mammalian iGluRs interact with residues in the LBD, with some structural overlap with binding sites for natural ligands (reviewed in [4]), we explored whether the three amino acid changes that converted the ligand specificity of GluE1αA from glycine to glutamate also altered sensitivity to AMPA, Kainate, NMDA, and quisqualate. Perfusing different extracellular solutions over recorded cells expressing the wildtype receptor revealed robust currents elicited by 1 mM glycine, but insensitivity to 1 mM AMPA, Kainate, and NMDA (Figure 4b and b). Similarly, the m2 variant of GluE1αA produced robust responses to 1 mM glutamate but was insensitive to the tested agonists. In contrast, the m3 variant exhibited moderate responses to 1 mM AMPA, with the drug causing slow desensitizing currents with peak amplitudes roughly 36% of those elicited by 1 mM glutamate (Figure 4a and b). We also tested the competitive AMPA/Kainate receptor blocker CNQX, uncovering a low affinity block of glycine-activated currents for the wildtype receptor with an IC_50_ of 210 ±60 µM. Interestingly, the m2 variant of GluE1αA exhibited a biphasic response, with 0.1 µM CNQX causing a near doubling of peak current amplitude (EC_50_ = 0.08 ±0.03 µM for a biphasic curve fit over the data), followed by potent block of glutamate-activated currents with concentrations >1 µM (IC_50_ = 0.96 ±0.07 µM for a monophasic curve fit over the blocked current, and 0.84 ±0.35 µM for a biphasic curve fit; Figure 4c to e). Instead, the m3 variant reverted to a monophasic sensitivity to CNQX, albeit with much higher affinity than the wildtype receptor with an IC_50_ of 5.05 ±2.45 µM (Figure 4c and d). Altogether, and perhaps as expected, mutations that alter ligand specificity can also alter sensitivity to pharmacological compounds that associate with the ligand binding site of iGluRs.

**Fig. 4:**
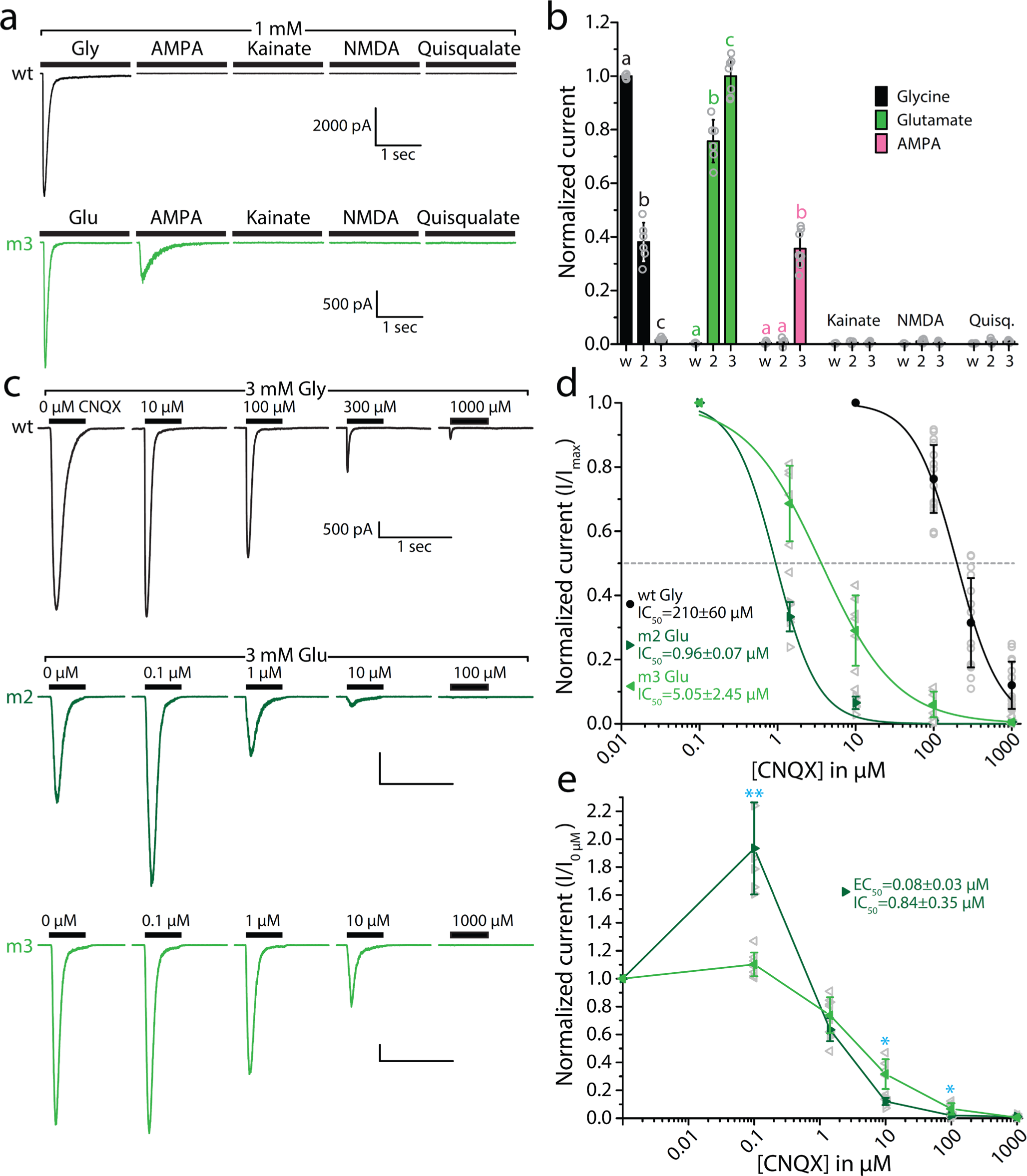
GluE1αA mutations that alter ligand specificity also affect pharmacological sensitivity to CNQX and AMPA. **a** Sample whole-cell recordings demonstrating that GluE1αA is not sensitive to 1 mM AMPA, kainate, NMDA, or quisqualate, wile the m3 variant is sensitive to AMPA. **b** Bar plot of average normalized peak inward currents of wt, m2, and m3 variants of GluE1αA in response to different agonists at 1 mM (n=6-7). Letters above the bars denote statistically significant differences among applied compounds for post hoc Tukey tests (p<1E-4) after One Way ANOVAs (glycine: F=809.23, p=5.55E-16; glutamate: F=367.97, p=1.79E-13; AMPA: F=126.50, p=4.07E-10). **c** Sample GluE1αA wt, m2 and m3 currents elicited by 3 mM glycine or glutamate with increasing concentrations of co-applied 6-cyano-7-nitroquinoxaline-2,3-dione (CNQX). **d** Dose response curve plot of average peak GluE1αA wt, m2, and m3 currents blocked by co-applied CNQX (n=7-15). One Way ANOVA confirmed significant differences of the IC_50_ values between wt and the two mutant variants (F=79.25, p=5.02E-12; p<1E-14 for Tukey post hoc tests), while the m2 and m3 variants were not different. Nonetheless, a two-sample T-test between mean m2 and m3 IC_50_ values was significant producing a p value of 1.56E-3. **e** CNQX dose response curves for the m2 and m3 variants of GluE1αA normalized to peak current amplitude in the absence of CNQX. EC_50_ and IC_50_ values obtained by fitting the m2 dose response data with a biphasic dose response curve [80], reflecting agonism then block by CNQX with increasing concentration, are indicated. The cyan asterisks above select data points indicate statistically significant differences between mean normalized current values of the m3 and m3 variants at different CNQX concentrations, as determined via two-sample T-tests (two asterisks: p<0.005; one asterisk: p<0.05).

### A unique serine residue in the NQR site decreases Ca*^2+^* selectivity and diminishes voltage-dependent regulation by polyamines

Interestingly, dynamic sequence changes in the NQR site of placozoan group 1 epsilon receptors (Figure 1b) suggests dynamic evolution of ion selectivity and polyamine regulation in this clade. In mammalian AMPA, Kainate, and NMDA receptors, the NQR site forms the narrowest part of the ion permeation pathway and is locus for regulating ion permeation and selectivity. Heteromeric AMPA and Kainate receptors for example, which encode glutamine in the NQR site at the mRNA level, exhibit different Ca^2+^ permeation properties depending on subunit composition, where A to I mRNA editing of select subunits converts the NQR glutamine to arginine, rendering channels that are impermeable to Ca^2+^ (reviewed in [3, 33]). NMDA receptors do not exhibit A to I editing, but an asparagine to lysine mutation in the otherwise invariable NQR site of the NMDA2A subunit is associated with severe developmental delay and early-onset epileptic encephalopathy, and *in vitro* this mutation causes severely diminished Ca^2+^ permeation and voltage-dependent Mg^2+^ block of conducted currents [40]. Interestingly, while all placozoan AKDF receptors, and all clade 2 and 3 Epsilon receptors exhibit glutamine resides in the NQR site, thus resembling mammalian AMPA and Kainate receptors, all GluE1 subunits possess divergent residues of either alanine, leucine, or serine (Figure 1b). Hence, to determine if the non-canonical serine residue in the GluE1αA NQR site impacts ion permeation and/or selectivity, as well as polyamine regulation, we characterized its ion permeation properties. First, we conducted bi-ionic reversal potential experiments with an invariable internal solution containing 150 mM Na^+^ and used perfusion to switch the extracellular solution from equimolar Na^+^ to Li^+^, K^+^, and Cs^+^ (Figure 5a). Eliciting currents with 3 mM glycine while changing the holding voltage to between -80 and +80 mV produced currents with reversal potentials (*i.e.*, E_Rev_ values) near zero mV for all bi-ionic conditions (Figure 5b to e). For the wildtype receptor, converting differences in E_Rev_ values for the different bi-ionic conditions to pX^+^/pNa^+^ permeability ratios (where X = Na, Li, K, or Cs) revealed a slight preference for Na^+^ over Li^+^ and Cs^+^ ions, and a slight preference for K^+^ over Na^+^ (Figure 5f; statistical comparisons provided in Table S4a). Mutation of the NQR residue to glutamine (S_642_Q), to resemble other placozoan iGluRs and AMPA/Kainate receptors, abrogated the marginal permeability differences of the wildtype receptor, with all pX^+^/pNa^+^ values being statistically indistinguishable from each other. Similarly, mutation to asparagine (S_642_N), to resemble NMDA receptors, also abrogated selectivity preferences except for Li^+^ which retained its marginally lower permeability compared to Na^+^. We also mutated the NQR site to arginine, resulting in non-selectivity between Na^+^ and Li^+^, and slightly diminished permeability of K^+^ and Cs^+^ relative to Na^+^ (Figure 5f). Altogether, it appears as though the serine in the NQR site of GluE1αA renders a channel that is largely non-selective between monovalent cations, similar to AMPA, Kainite, and NMDA receptors.

**Fig. 5:**
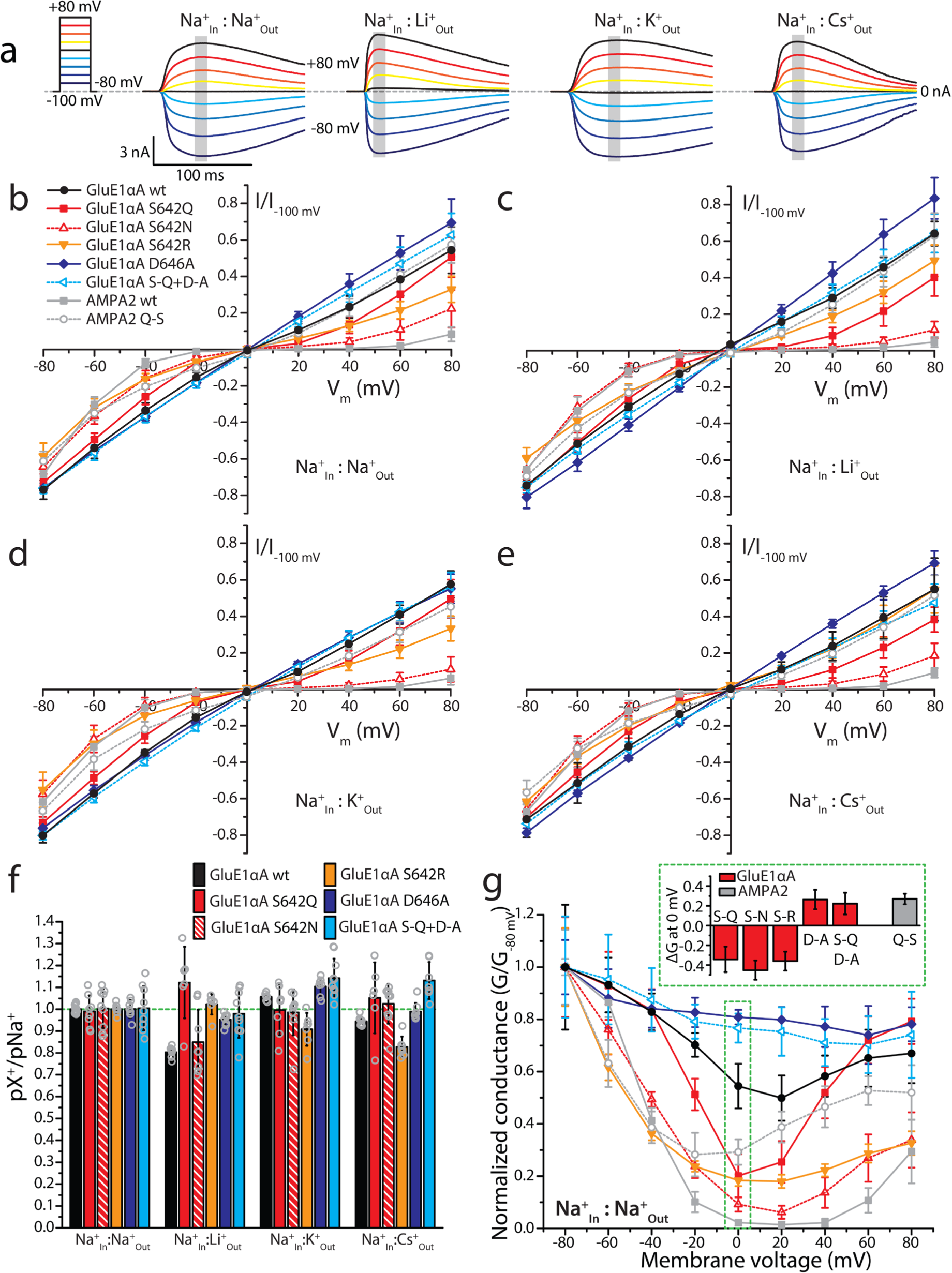
A non-canonical serine in the QRN site of GluE1αA contributes to non-selective monovalent currents and weak rectification indicative of reduced polyamine block. **a** Sample wildtype GluE1αA whole-cell currents elicited by 3 mM glycine recorded at membrane voltages ranging from -80 mV to +80 mV under bi-ionic conditions of 150 mM intracellular Na^+^ and 150 mM extracellular X^+^ ions where X is Na, Li, K or Cs. **b** Plot of average normalized current vs. membrane voltages (I-V) under equimolar intracellular and extracellular Na^+^ conducted by wildtype and pore mutant variants of GluE1αA, the wildtype human AMPA2 receptor, and a mutant AMPA2 variant bearing a glutamine (Q) to serine (S) mutation in the pore QRN site (n=6-13). **c** Similar I-V plot as panel b but with equimolar internal Na^+^ and external Li^+^ (n=6-9). **d** I-V graph with equimolar internal Na^+^ and external K^+^ (n=7-10). **e** I-V plot with equimolar internal Na^+^ and external Cs^+^ (n=6-9). **f** Average permeability ratios (pX^+^/pNa^+^) ± standard deviation calculated from the I-V data presented in panels b to e. **g** Average normalized conductance vs. membrane voltage (G-V) plot derived from the I-V data in panel b for the different receptors and variants under symmetric bi-ionic Na^+^ conditions. **Inset:** plot of differences in the normalized conductance data for all receptor variants relative to their wildtype counter parts at 0 mV (*i.e.*, values demarked by the green dashed box in the main plot). Differences between average wildtype and corresponding variant conductance data at 0 mV were all significant in paired T-tests (p≤5.5E-6).

Despite subtle differences in monovalent ion selectivity, we did notice differences in the rectification of currents through changing voltage for the different pore variants. Specifically, while the wildtype receptor exhibited only slight rectification with an almost linear current-voltage (I-V) relationship, the S_642_N, S_642_Q, and S_642_R variants showed increased current rectification near 0 mV (Figure 5b to e). Converting the I-V plots to conductance-voltage (G-V) plots, which removes the effect of driving force and elucidates changes in conductance as a function of voltage, revealed a continued decline in conductance for the wildtype channel with increasing depolarization up to +20 mV, followed by a subsequent increase at more positive voltages (Figure 5g). This voltage-dependent drop in conductance, also apparent for the *in vitro* expressed Epsilon receptors from the cephalochordate *B. lanceolatum* and the ctenophore *M. leidyi* [5, 8], is consistent with a mild voltage-dependent block of GluE1αA by polyamines. These compounds, namely spermine, spermidine, and putrescine, are endogenously expressed in the cytoplasm of all living cells [41], where they regulate various targets including AMPA, Kainate, and Delta receptors [11, 42, 43]. Indeed, previous studies have demonstrated that polyamine block depends on NQR residues, as well as a conserved aspartate 4 amino acids downstream (*i.e.*, the +4 site) [11]. These linear cationic molecules are thought to insert into the pore selectivity filter at depolarized voltages, facilitated by electrostatic interactions with NQR and +4 aspartate residues, resulting in obstructed ion permeation [44]. As noted earlier, most placozoan iGluRs, including GluE1αA, have a conserved acidic (*i.e.*, aspartate or glutamate) residue in the +4 site (Figure 1a), also apparent in most metazoan epsilon and AKDF receptors, as well as some cnidarian NMDA receptors [5].

Notably, although current rectification and a corresponding drop in conductance was apparent for the wildtype GluE1αA receptor, it became much more pronounced in the AMPA/Kainate/NMDA-like variants bearing glutamine, asparagine, or arginine in the NQR site (Figure 5b to e, g and Supplementary Figure 7). Indeed, the difference in normalized conductance values at 0 mV between the wildtype receptor and the S_642_Q, S_642_N, and S_642_R variants revealed negative changes ranging from 34.3 ±12.9% to 45.3 ±9.8%, reflecting considerable increases in polyamine block of the variant channels (Figure 5g inset). We also mutated the +4 aspartate GluE1αA to alanine (D_646_A), causing only marginal effects on monovalent ion selectivity, with a slight but statistically significant increase in K^+^ over Na^+^ permeability (Figure 5f). However, this mutation resulted in completely linear I-V and G-V curves (Figure 5b to e, g and Supplementary Figure 7), and a positive change in normalized conductance at 0 mV relative to wildtype (*i.e.*, 26.4 ±9.8%; Figure 5g inset), indicating a pronounced loss of polyamine block. Similar observations were made for a double mutant bearing the S_642_Q mutation, which on its own increased polyamine block, with the D_646_A mutation, pointing to a conserved and dominant function of the +4 residue in polyamine regulation.

To determine whether the observations made for the *T. adhaernes* Epsilon receptor are relevant in a broader context, we conducted similar experiments on human AMPA2 bearing either a wildtype NQR glutamine or a GluE1αA-like serine. Interestingly, while the wildtype receptor exhibited characteristic rectifying currents and pronounced U-shaped G-V curves under all ionic conductions (activated by 3 mM glutamate), the serine variant had much more linear I-V curves and diminished drops in conductance with increasing depolarization (Figure 5b to e, g and Supplementary Figure 7). This mutation did not cause appreciable changes in current reversal potentials, although the strong rectification of the wildtype receptor prevented us from accurately determining E_Rev_ values for making statistical comparisons. Nonetheless, it seems that a serine residue in the AMPA2 NQR site does not impose strong changes in monovalent ion selectivity, as demonstrated for GluE1αA. Lastly, comparing the difference in normalized conductance at 0 mV between the wildtype AMPA receptor and the serine variant revealed a positive change in normalized conductance of 27.0 ±5.4%, consistent with reduced polyamine block. Altogether, our data indicate that the presence of a serine residue in the NQR site diminishes but does not abrogate polyamine regulation, and furthermore, that the acidic +4 aspartate in Epsilon iGluRs plays a conserved and critical function in polyamine regulation, similar to AMPA, Kainate, and perhaps other receptor types.

Since the NQR site also impacts Ca^2+^ permeation, we conducted bi-ionic electrophysiology experiments using an external solution of 4 mM Ca^2+^ and different internal solutions bearing either Na^+^, Li^+^, K^+^, or Cs^+^ at 100 mM. Eliciting currents with 3 mM glycine through voltage steps between -100 and +80 mV produced large outward currents for the wildtype GluE1αA receptor, compared to comparatively smaller inward Ca^2+^ currents under all bi-ionic conditions (Figure 6a). Currents for the wildtype receptor exhibited negative reversal potential values, ranging between -59.2 ±4.5 mV for K^+^_In_:Ca^2+^_Out_ and -46.9 ±2.0 mV Cs^+^_In_:Ca^2+^_Out_ (Figure 6b to e). Converting E_Rev_ values to pCa^2+^/pX^+^ values (where X = Na, Li, K, or Cs) revealed only slight changes in permeability ratios between the four conditions ranging between 0.67 ±0.11 (for K^+^_In_:Ca^2+^_Out_) to 1.14 ±0.10 (for Cs^+^_In_:Ca^2+^_Out_) (Figure 6g and Supplementary Figure 8a; statistical comparisons provided in Table S4b). We made similar recordings of the wildtype AMPA receptor, finding considerably right-shifted I-V curves, with strong rectification of currents near the reversal preventing us from obtaining accurate E_Rev_ values (Figure 6b to e and Supplementary Figure 8a). Notably different between the human and *T. adhaerens* receptors is the ratio of inward Ca^2+^ current at -100 mV vs. outward monovalent current at +80 mV. Specifically, while the inward Ca^2+^ current ranged between 26.3 ±5.6% and 50.6 ±8.8% of the outward monovalent current for GluE1αA (for the conditions K^+^_In_:Ca^2+^_Out_ and Cs^+^_In_:Ca^2+^_Out_, respectively), the relative inward Ca^2+^ current for AMPA2 ranged between 175.1 ±25.3% for Na^+^_In_:Ca^2+^_Out_, and 892.1 ±107.0% for Li^+^_In_:Ca^2+^_Out_;, the latter attributable to very small amplitude outward currents when Li^+^ was present as the outward-permeating ion (Figure 6b to e and Supplementary Figure 8a). In G-V plots normalized to -80 mV, a difference in inward Ca^2+^ permeation is also apparent between GluE1αA and AMPA2. That is, while the normalized inward Ca^2+^ conductances of AMPA2 at -80 mV range between 77.4 ±5.2% (Na^+^_In_:Ca^2+^_Out_) and 382.4 ±9.7% (Li^+^_In_:Ca^2+^_Out_), those for GluE1αA were much lower ranging between 27.9 ±4.5% (K^+^_In_:Ca^2+^_Out_) and 49.6 ±5.7% (Cs^+^_In_:Ca^2+^_Out_; Figure 6g and Supplementary Figure 8b to e). Altogether, these observations indicate that the *T. adhaerens* GluE1αA receptor bearing a serine in the NQR site is less permeable to Ca^2+^ compared to AMPA2.

**Fig. 6.**
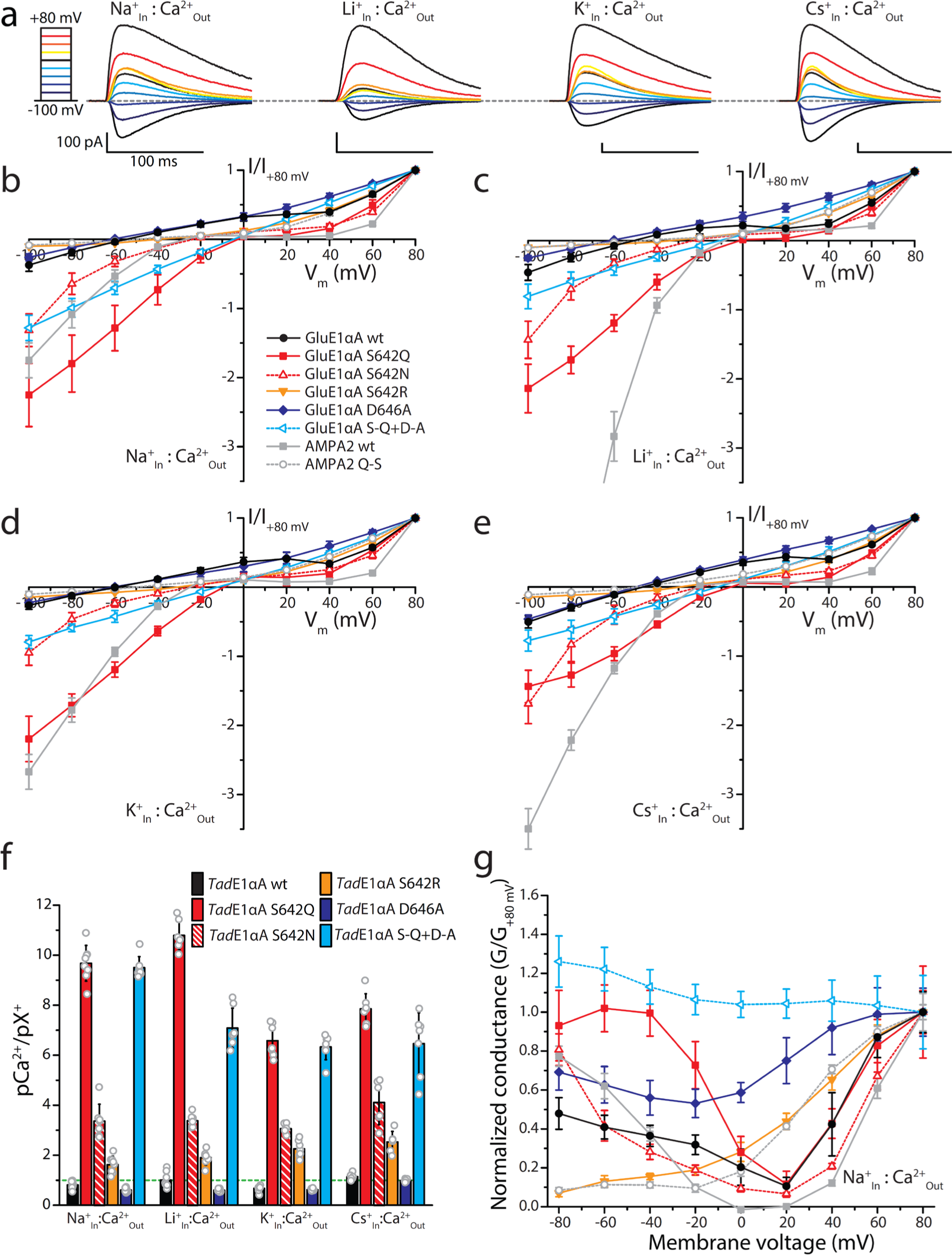
The NQR serine renders reduced Ca^2+^ permeation for GluE1αA compared to variants with canonical Q, N, and R residues. **a** Sample wildtype GlueE1α whole-cell currents recorded at membrane voltages ranging from -100 to +80 mV under bi-ionic conditions of 4 mM extracellular Ca^2+^ and 100 mM intracellular X^+^ where X denotes Na, Li, K or Cs. **b** Plot of average normalized current vs. membrane voltages (I-V) under 4 mM extracellular Ca^2+^ and 100 mM intracellular Na^+^ conducted by wildtype and pore mutant variants of GluE1αA, the wildtype human AMPA2 receptor, and a mutant AMPA2 variant bearing a glutamine to serine mutation in the QRN site (n=7-11). **c** Similar I-V plot as in panel b but with 100 mM internal Li^+^ instead of Na^+^ (n=7-8). **d** I-V plot with 100 mM internal K^+^ (n=7-8). **e** I-V plot with 100 mM internal Cs^+^ (n=6-8). **f** Average permeability ratios (pCa^2+^/pX^+^) ± standard deviation calculated from the I-V data presented in panels b to e. **g** Average normalized conductance vs. membrane voltage (G-V) plot derived from the I-V data in panel b for the different receptors and variants under external Ca^2+^ and internal Na^+^ conditions.

To test whether a serine residue in the NQR site of AMPA2 imparts similar low Ca^2+^ permeation, we conducted parallel experiments with the serine variant. Despite our inability to calculate accurate reversal potentials for this variant due to its strong rectifying currents, we observed clear leftward shifts in the I-V curves under all bi-ionic conditions, along with marked reductions in the ratio of inward Ca^2+^ current at -100 mV relative to outward monovalent current at +80 mV (Figure 6b to e), concurrent with reductions in inward Ca^2+^ conductance at negative voltages in the G-V plots (Figure 6g and Supplementary Figure 8b to e). Thus, a serine residue in the NQR site significantly reduces Ca^2+^ vs. monovalent permeability and Ca^2+^ conductance in AMPA2 receptors.

Next, we tested whether the AMPA/Kainate/NMDA-like NQR variants of GluE1αA exhibit enhanced Ca^2+^ permeation properties. In agreement with expectations, both glutamine and asparagine in the NQR site led to strong rightward shifts in the I-V curves under all bi-ionic conditions (Figure 6b to e), and corresponding strong increases in pCa^2+^/pX^+^ ratios ranging between 6.6 ±0.6 (Cs^+^_In_:Ca^2+^_Out_) and 10.8 ±0.5 (Li^+^_In_:Ca^2+^_Out_) for the S_642_Q variant, and between 3.1 ±0.2 (K^+^_In_:Ca^2+^_Out_) and 4.1±0.8 (Cs^+^ :Ca^2+^_Out_) (Figure 6f). There were also marked increases in normalized inward Ca^2+^ conductance at negative voltages (Figure 6g and Supplementary Figure 8b to e), as well as U-shapes G-V curves consistent with increased polyamine block. Altogether, this data supports a significant increase in Ca^2+^ permeability and conductance when the NQR serine is mutated to resemble mammalian iGluRs. In contrast, the S_642_R mutation, despite causing mild rightward shifts in I-V plot reversal potentials and mild increases in pCa^2+^/pX^+^ values, produced very small inward Ca^2+^ currents at -100 mV relative to outward monovalent currents at +80 mV (Figure 6b to e), and significantly diminished normalized inward Ca^2+^ conductance at negative voltages (Figure 6g and Supplementary Figure 8b to e). These observations are consistent with those made for homotetrameric mammalian AMPA and Kainate receptors bearing four arginine residues in the NQR sites, which also became poorly conductive to cations [45], perhaps due to repulsion of permeating cations by the positively-charged arginine side chains [46].

Lastly, we observed only minimal changes imposed by the D_646_A mutation, despite a complete loss of an obvious kink in the I-V curves presumably caused by polyamine block (Figure 6b to e), also apparent in the near constant normalized G-V plots (Figure 6g and Supplementary Figure 8b to e). This variant had similar albeit statistically lower pCa^2+^/pX^+^ values compared to wildtype ranging between 0.63 ±0.04 (Li^+^ :Ca^2+^) and 1.04 ±0.06 (Cs^+^_In_:Ca^2+^_Out_), and increased normalized conductances across all voltages, likely due to loss of polyamine block (Figure 6g and Supplementary Figure 8b to e). Combining the S_642_Q and D_646_A mutations yielded increases in pCa^2+^/pX^+^ ratios relative to wildtype, similar to S_642_Q on its own, except in the presence of Li^+^ where the single mutant had a value of 10.79 ±0.54 compared to only 6.26 ±0.66 for the double mutant (Figure 6f). Altogether, these experiments corroborate our previous observations that a serine residue in the NQR site diminishes polyamine block, and in addition, reduces Ca^2+^ permeation. Furthermore, the +4 aspartate residue also plays a central role in polyamine regulation in the presence of Ca^2+^ ions.

### Insights into polyamine regulation of GluE1αA by modelling and extracellular perfusion of polyamines

To gain more insights into the regulation of GluE1αA by polyamines, we used homology modelling and docking to explore the energy properties of spermine in the pore of the wildtype receptor bearing a serine residue in the NQR site, and the asparagine S_642_N variant which exhibited significantly enhanced polyamine block (Figures 5 and 6). To start, we used AlphaFold 2 to predict the tetrameric structure of both variants lacking their N-terminal domains (i.e., V_416_ to I_921_ of NCBI accession number PP886186). Both predicted structures resembled a closed pore configuration, perhaps expected given that most experimental structures used for training the AlphaFold artificial neuronal network were in the closed configuration. Because we were interested in exploring how spermine block occurs in the open, conducting state, we modelled the open-pore domain for our spermine docking experiments. To build such models, we isolated the pore domain of the predicted GluE1αA structures (A_554_ to V_682_ and K_833_ to I_921_ of PP886186) and 3D-aligned these to the cryo-EM structure of the AMPA2 receptor in the open state (PDB number: 6o9g; [44]). This was achieved by minimizing root mean square (RMS) deviations of alpha carbon atoms in sequentially matching positions of the first poor-loop helix of the M2 domains of the two receptors (as reported previously [47]). To reduce misalignment of key P-loop residues in the NQR and +4 site, we then Monte Carlo-minimized the RMS distances between matching alpha carbons in the M1 and M3 of the AlphaFold models and the AMPA2 cryo-EM structure. Finally, we used the in silico opened models of the wildtype and D_646_A variants of GluE1αA to predict energetically optimal binding modes of spermine in the pore (Figure 7a to d). This was done by using the ZMM GRID module (see Materials and Methods) to pull spermine from the cytoplasm to the extracellular side of each GluE1αA variant structure with steps of 0.5 angstroms, and Monte Carlo energy minimization of the structure at each step. Translation of the spermine central atom along the pore axis (its z-coordinate) was frozen, backbone alpha carbon atoms were pinned, and all other generalized coordinates of the system were free to move during energy minimizations (Supplementary Figure 9a and b). Energy plots of spermine-receptor interactions reveal declining electrostatic (ELRT) and non-bonded (ELRN) energy components for both the wildtype and S_642_N variants as spermine enters the pore, while the desolvation energy (ELRD) increases reflecting increasing dehydration. Despite the latter, both receptors had decreasing total energy values (ELRT) as spermine moved into the pore, with minima (and hence most favourable) positions at step 32 for the wildtype receptor and step 34 for the S_642_N variant (Supplementary Figure 9b and c). In the wildtype receptor, the cytoplasmic/lower amino group of spermine forms strong hydrogen bonds with the +4 D_646_ residues of subunits 3 and 4 (Figure 7a and b), which contribute the lowest energies of the receptor-ligand interactions, while the same aspartates from subunits 1 and 2 also contribute (Table S3d and e). Similarly, all four D_646_ residues in the S_642_N variant contribute low energies for interactions with spermine, albeit with small amplitudes than wildtype (Figure 9c and d; Table S3d to e). These observations are overall consistent with our experimental data where D_646_A variants of GluE1αA exhibited a near complete loss of polyamine block evident as negligible current rectification (Figures 5 and 6). Also shared between both structures are predicted electrostatic interactions between upper/central amino group in the spermine molecule and highly conserved glutamine residues one position downstream of the NQR site (Figure 1b), in both cases involving subunits 1 and 4 (Figure 7a to d). Also consistent with our experimental data is the more negative total energy scores of spermine in step/position 34 of the S_642_N variant, compared to the wildtype receptor at step 32, the former having a total receptor-ligand score of -20.32 kcal/mol, and the latter of -16.38 kcal/mol (Table S3d). In part, this marked energy difference is attributable to the NQR asparagine residues which collectively contribute -6.32 kcal/mol to the total spermine-receptor energy, forming numerous predicted hydrogen bonds with the extracellular/upper ammonium group of spermine, while the NQR serine residue in subunit 1 of the wildtype receptor contributes only -0.78 kcal/mol, and the same serine on subunit 3 provides a net repulsive energy of 0.42 kcal/mol (Table S3d).

**Fig. 7.**
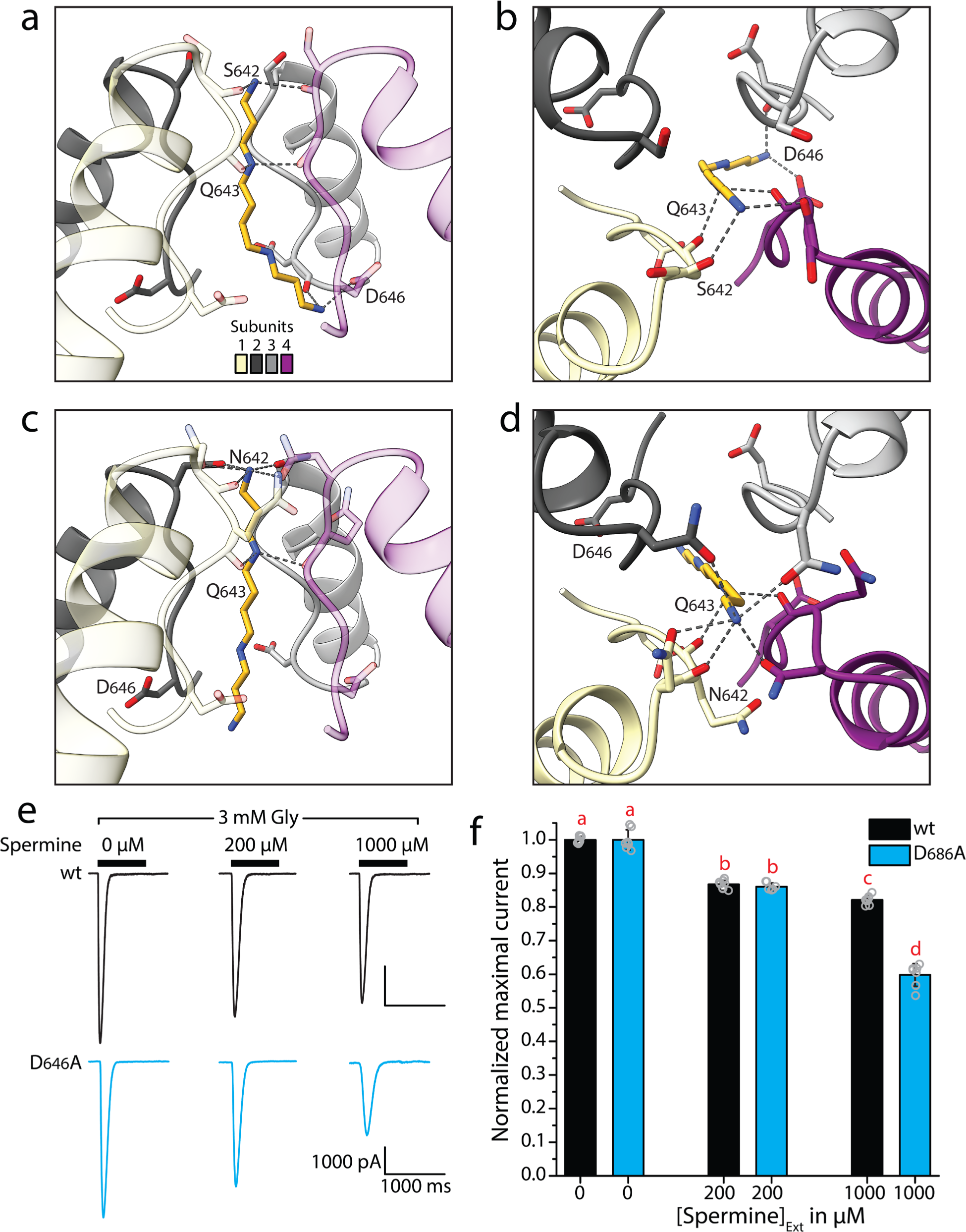
GluE1αA pore residues contribute to intracellular and extracellular regulation by polyamines. **a** Homology model and Monte Carlo-minimized structure of spermine in the open GluE1αA pore domain (side view, step 32). **b** Top-down view of the structure in panel a. **c** Homology model and Monte Carlo-minimized structure of spermine in the open pore domain of the S_642_N variant of GluE1αA (step 34). **d** Top-down view of the structure in panel c. In panels a to d, predicted hydrogen bonds between pore-loop residues of the subunits and spermine are depicted by dashed lines. **e** Sample wildtype and D_646_A currents elicited with glycine co-applied with increasing concentrations of perfused (extracellular) spermine. **f** Bar plot of normalized currents elicited by 3 mM glycine with and without co-applied spermine. Letters above the bars denote statistically significant differences among and between conditions determined by post hoc Tukey tests (p<0.005) after One Way ANOVAs (F=385.66, p<1E-14).

As a final set of experiments, we used perfusion to determine whether spermine applied extracellularly causes changes to ligand-induced macroscopic currents of GluE1αA, as has been reported for select mammalian iGluRs (reviewed in [3]). Application of 200 µM spermine caused a modest reduction in peak inward current of 13.3 ±1.3% relative to currents elicited by 3 mM glycine alone, while 1000 µM spermine reduced peak current by 17.9 ±1.3% (Figure 7e and f). Since this observed block could be due to spermine molecules entering the pore from the extracellular surface, we also tested the D_646_A variant, reasoning that this variant should have altered sensitivity if the observed effects are due to extracellular polyamines entering the pore. Indeed, 1000 µM spermine caused a more pronounced block for the D_464_A mutant, reducing peak current by 40.2 ±3.4%, while block caused by 200 µM spermine was indistinguishable from that of the wildtype receptor (*i.e.*, 14.0 ±1.1%; Figure 7e and f). Although the reason for this is unclear, it is notable that neutralization of the +4 aspartate residues via alanine substitution removes a ring of negatively charged carboxyl side groups which are thought to facilitate the transfer of polyamines across the pore, whereby extracellular polyamines moving through the receptor could become trapped in upper regions of the pore, unable to effectively progress into the cytoplasm and hence block inward current.

## Discussion

### An updated phylogeny of placozoan iGluRs

Our phylogenetic analysis is consistent with several recent studies and provides several new insights, particularly into the evolution of iGluRs within the phylum Placozoa. One notable insight is the apparent dichotomy between AKDF and Epsilon receptors, the former exhibiting one-to-one orthology among the four examined species, despite their deep branch separation (Figure 1b). Also similar between species is their enriched expression in digestive lipophil cells. Epsilon receptors on the other hand appear to have undergone lineage-specific duplications and losses, resulting in much more phylogenetic diversity, combined with more heterogeneity in cellular expression across species (Figure 1b). Consistent with the species phylogeny [24], *T. adhaerens* and *Trichoplax* species H2 share more orthologous receptors with each other, as do *H. hongkongensis* and *C. collaboinventa*, and these are often expressed in the same cell type. Of note, during our manual curation and analysis of placozoan iGluRs we identified a small number of Epsilon receptor sequences from each species that were too fragmented to include in our phylogenetic analysis. Indeed, as more gene data becomes available, for these or perhaps other placozoan species, more complete iGluR sequences might emerge that will help better resolve the phylogeny of placozoan Epsilon receptors. Nonetheless, the three strongly supported clades of Epsilon receptors delineated in our analysis will likely withstand the addition of new sequences, which by extension, can be named according to our proposed cladistic naming scheme.

Interestingly, like AKDF receptors, the GluE3α and GluE3β receptors show enriched expression in lipophil cells, while GluE1 receptors tend to be enriched in neuron-like peptidergic cells (Figure 1b). Currently, our knowledge on the physiological functions of iGluRs in non-bilaterian animals is very limited. In cnidarians, glutamate is not considered to be a major transmitter used in synapses, although it elicits various behavioral responses in select species, mainly associated with feeding including chemotaxis, discharge of cnidocytes used for hunting and defense, and muscle contractions (reviewed in [1]). Indeed, the enriched expression of iGluRs in placozoan lipophil cells, which are involved in feeding, might indicate a similar function in placozoans, and perhaps more broadly, that a primordial function of iGluRs, before the evolution of synapses, was for sensing amino acids in the environment to identify food sources and mediate feeding behaviors. Instead, the expression of iGluRs in neuron-like peptidergic cells might reflect roles more akin to neural signaling, where for example the receptors could serve to depolarize cells in response to ligands present in the environment, triggering secretion of peptides that regulate behavior. As noted earlier, placozoans elicit numerous behavioral responses to applied transmitters including glycine, glutamate, and GABA [14, 15]. However, like cnidarians, there is no evidence suggesting that placozoans actively secrete these substances into the environment in a regulated manner to affect other cells or coordinate behavioral responses, although some cells are reported to possess pale vesicles, in addition to dense core, the former consistent with small molecule transmitters [48, 49]. Although possible, something that needs to be reconciled is how effective extracellular concentrations are reached, given that placozoans lack synapses [49], or tight/septate junctions between epithelial cells that would prevent secreted molecules from quickly diffusing out of the animal interior [50, 51].

Prior to this study, the only placozoan iGluR to be functionally characterized *in vitro* is the AKDF1 receptor from *T. adhaerens* [30]. As noted, this receptor lacks an aspartate residue in its ligand binding domain that in other receptors interacts with the backbone amino group of amino acid ligands (D_705_), leading to constitutive leak currents that are blocked by various ligands, most prominently glycine as predicted by its ligand-binding sequences. This same aspartate to tyrosine adaptation is evident in all other placozoan AKDF1 orthologues (Figure 1b), indicating this is an ancestral adaptation within the phylum. Interestingly, several Epsilon receptors also lack ligand backbone interacting residues, including members of the GluE1 clade which also lack D_705_ residues. These same receptors also have sequence differences in their ligand binding domains that prevent prediction of their ligand specificity. As such, this clade of receptors might have unique ligand binding properties. Lastly, it is evident that evolutionary changes in ligand selectivity are apparent within both the AKDF and Epsilon receptors in placozoans, consistent with predictions of ligand specificity changes among the ctenophore Epsilon receptors [8] and more broadly within across all major lineages of metazoan iGluRs [10].

### Insights into iGluR ligand-specificity

Our electrophysiological characterization of GluE1αA revealed a broad ligand selectivity, with sub-millimolar sensitivity to glycine, alanine, and D-/L-serine, and millimolar sensitivity to valine (Figure 2). In its preference for glycine over glutamate, GluE1αA resembles other homomeric Epsilon receptors from *B. lanceolatum* (GluE1) and *M. leidyi* (ML032222a) but differs from a second *M. leidyi* receptor (ML05909) that is activated by both glutamate and glycine [5, 8]. Notable is that alanine was the most potent activator of GluE1αA, a hydrophobic amino acid that also activates the glycine-sensitive NMDA1/GluN1 subunit [52], as well as the divergent GluR1 receptor from the rotifer *Adineta vaga*, a highly atypical receptor that resembles bacterial iGluRs phylogenetically, structurally, and in its selectivity for K^+^ ions over Na^+^ [53, 54]. This receptor, which is also activated by glutamate but not glycine, is more broadly sensitive to hydrophobic amino acids compared to GluE1αA and GluN1, exhibiting strong activation by methionine, cysteine, and moderate activation by phenylalanine [54]. Interestingly, X-ray structures and functional studies uncovered unique ligand properties of the *A. vaga* GluR1 receptor, whereby chloride ions act as co-activators for hydrophobic amino acids by forming surrogate interactions with residues in the ligand binding domain that normally associate with the glutamate side chain [54]. Since the wildtype *T. adhaerens* receptor did not respond to glutamate, while the m3 variant responded to glutamate but not alanine (Figure 3b), it seems unlikely a similar mechanism accounts for the activation of GluE1αA by alanine and the also hydrophobic amino acid valine (Figure 2b and c).

By mutating just three amino acids in the ligand binding domain, we were able to completely switch the ligand selectivity of GluE1αA from glycine/serine to glutamate, providing experimental support for the hypothesis that just a small set of residues in the ligand binding domain determine ligand specificity. In fact, the m2 variant bearing just two mutations (S_653_G and I_655_T) changed the selectivity mostly towards glutamate, while the third mutation, F_732_Y, increased sensitivity to glutamate (Figure 1d), almost completely attenuated glycine sensitivity (Figure 1b and c) and caused nascent sensitivity to AMPA (Figure 4a and b). In our models, this mutated tyrosine residue forms salt bridge with the reside D_705_, which in the wildtype channel forms key contacts with glycine, leading to reduced binding energy between D_705_ and glycine in the mutant receptor (Table S3). Perhaps, this contributes to the near complete loss of glycine sensitivity of the m3 triple mutant. In addition, the F_732_Y mutation would introduce a hydroxyl group into the ligand binding pocket, increasing its polarity and perhaps helping to attract the side chain carboxyl group of glutamate into the pocket. According to the ligand selectivity hypothesis, when position 655 in the ligand binding domain is hydrophobic, there is a presence for glycine, while a polar threonine contributes to glutamate selectivity [8, 10]. In wildtype GluE1αA, this equivalent position bears an isoleucine, which we mutated to a threonine in the m2 and m3 variants (Figure 3a). Like the F_732_Y mutation, the I_655_T mutation would introduce a hydroxyl group into ligand pocket increasing its polarity, and in our models, this residue forms hydrogen bonds with the glutamate side chain (Figure 3i and Supplementary Figure 6d; Table S3). Additionally, mutation of serine at position 653 to glycine would enlarge the binding pocket, perhaps better accommodating the larger glutamate side chain (Figure 3h and I and Supplementary Figure 6c and d).

Interestingly, the three mutations that switched the ligand selectivity of GluE1αA from glycine to glutamate also disrupted sensitivity to alanine, L-and D-serine, and valine, making the receptor more similar to AMPA and Kainate receptors that are more exclusively gated by glutamate over other amino acids (reviewed in [3]). If this observation extrapolates as a general feature of iGluRs, it may point to one of the selective pressures that drove evolutionary changes in ligand selectivity, with conceivable advantages for detecting just glutamate in some contexts (*i.e.*, in synapses or the environment), or a broader set of amino acids in other contexts. Also interesting was that the mutations that switched ligand selectivity of GluE1αA also imposed nascent sensitivity to AMPA (Figure 4 and b) and increased sensitivity to the blocker CNQX (Figure 4c to e), further likening the placozoan receptor to vertebrate AMPA and Kainate receptors. Together, these observations suggest that structural features of the ligand binding domain that promote exclusive glutamate sensitivity also increase a receptor’s affinity for pharmacological compounds that are selective for such channels. However, there are apparently structural features that do not overlap for binding natural ligands vs. pharmacological compounds, since compounds that discriminate between the different vertebrate receptors do not necessarily do so for their invertebrate counterparts. For example, the heteromeric iGluRs that are found in the fruit fly neuromuscular junction, which are phylogenetically classified as Kainate receptors, are insensitive to Kainate [55], while the AMPA receptor GLR-1 from the worm *Caenorhabditis elegans* is activated by Kainate [56].

### Insights into ion permeation and polyamine regulation

Our phylogenetic analysis revealed that all placozoan GluE1 type receptors bear atypical NQR residues of serine, leucine, or alanine, while the remaining Epsilon receptors, and the four AKDF receptor types, bear ‘canonical’ glutamine residues also found in AMPA and Kainate receptors (Figure 1b). Similar variability in the NQR site is apparent among the Epsilon receptors from the ctenophore species *M. leidyi*, most bearing a glycine [8]. This variability extends to Epsilon receptors from cephalochordates, for example GluE1 from *B. lanceolatum* which bears a phenylalanine, and indeed all classes of metazoan iGluRs [5]. Perhaps the least variable among these are the AKDF receptors, that like the *T. adhaerens* AKDF receptors, as well as AMPA, and Kainite receptors, tend to have a glutamine in the NQR site [5]. Nonetheless, the general prevalence of glutamine in the NQR site of AKDF receptors, and its frequent occurrence in Epsilon receptors, suggests this residue was acquired through common descent. By extension, Epsilon receptors might have undergone evolutionary changes that relaxed selective pressures for retaining an NQR glutamine, allowing for diversification in ion conducting properties and regulation by polyamines.

Our characterization of GluE1αA revealed that the presence of a serine in the NQR site, which has a shorter side chain than glutamine and asparagine and bears a hydroxyl rather than an amino group, had minimal consequences on monovalent vs. monovalent selectivity (Figure 5 and Supplementary Figure 7). However, compared to glutamine and asparagine, the NQR serine significantly diminished Ca^2+^ permeation, and this was also apparent for the human AMPA2 receptor bearing a Q to S mutation in the NQR site (Figure 6 and Supplementary Figure 8). Conversely, mutating the GluE1αA NQR serine to glutamine and asparagine increased Ca^2+^ permeation by roughly 10-fold and 3-fold, respectively (Figure 6f), perhaps attributable to stronger coordination of Ca^2+^ ions facilitated by the longer side chains bearing carbonyl oxygen atoms.

Physiologically, it is notable that the GluE1 clade of Epsilon receptors, all with variable NQR sites, tend to be in peptidergic cells (Figure 1b). Based on single cell transcriptome analysis these resemble primordial neurons both in their expression of genes involved in neural development, and genes required for regulated secretion of signaling molecules including voltage-gated calcium channels and machinery for presynaptic exocytosis [24]. In these cells, expression of a receptor with an NQR serine would permit ligand-induced depolarization of the membrane voltage while diminishing the added effect of Ca^2+^ signaling, perhaps serving a functional purpose. Such an evolutionary change is not without precedent, occurring for example in voltage-gated sodium (Na_V_) channels. Here, ancestral Na_V_ channels that are enriched in aspartate/glutamate residues in the selectivity filter (*i.e.*, pore motifs of DEEA) are highly permeable to Ca^2+^, while bilaterian Na_V_1 channels evolved a lysine residue in domain III (*i.e.*, DEKA motifs) to become highly selective for Na^+^ over Ca^2+^. Interestingly, a parallel adaptation occurred in cnidarian Na_V_2 channels, but in domain II (*i.e.*, DKEA motifs), leading to convergent evolution of sodium selectivity [57].

Our experiments also indicate that an NQR serine, in both GluE1αA and AMPA2, diminishes but does not abrogate polyamine regulation. Instead, mutation of this residue in the *T. adhaerens* receptor to glutamine, asparagine, or lysine increased current rectification, most prominent for the S_642_N variant (Figures 5 and 6). According to our modeling, this higher degree of rectification is consistent with higher binding affinity of polyamines in the pore when asparagine is present compared to serine (Figure 7 and Supplementary Figure 9; Table S3). Interestingly, concurrent neutralization of the +4 aspartate to alanine (D_646_A) completely abrogated this enhanced polyamine block, while having minimal impacts on ion permeation, indicating this latter residue plays a key role in polyamine regulation. This was also reported for AMPA and Kainate receptors, where mutation of respective +4 aspartate and glutamate residues decreased current rectification and polyamine block while having minimal impacts on divalent ion permeation [58, 59]. As such, our data suggests that regulation of iGluRs via the NQR and +4 sites is an ancestral feature shared between AKDF and Epsilon receptors. Indeed, most placozoan receptors, except those from the clades GluE2β and GluE3γ, also bear acidic aspartate or glutamate residues at the +4 site (Figure 1b), and acidic residues are evident more broadly among all types of metazoan iGluRs, including cnidarian NMDA receptors and poriferan Lambda receptors [5]. Even plant iGluRs, most of which bear hydrophobic residues in the NQR site (*i.e.*, phenylalanine or tyrosine), have +4 glutamate residues [60], as does the primordial iGluR receptor from bacterial, GluR0, which bears a potassium channel-like selectivity filter but a +4 aspartate equivalent [61]. Whether conservation of this residue in non-metazoan receptors reflects even deeper origins for polyamine regulation is unclear, but it is notable that chimeric receptors comprised of Kainate receptors (GluR6) bearing the pore module of a plant iGluR (*i.e.*, AtGLR1.1 from *Arabidopsis thaliana*) conduct rectifying currents that are consistent with voltage-dependent regulation by polyamines [62]. Like resolved iGluR structures, the four D_646_ residues of the heteromeric GluE1αA receptor complex are predicted to come together at the cytoplasmic entrance of the pore, well positioned to interact with the positively charged amino groups of cytoplasmic polyamines. That this ring of aspartate residues provides an electrostatic entry point for polyamines to enter the pore was also indirectly supported by our perfusion experiments, where the D_646_A variant showed increased current block by 1 mM external polyamines compared to the wildtype receptor (Figure 7f). Here, we suggest that the absence of the aspartate residues in the mutant channel prevented extracellular polyamines from traversing the pore into the cell interior, hance accumulating inside the pore and blocking inward cation permeation.

In sum, it seems sequence changes in the pore-loop of iGluRs occurred quite commonly, representing adaptive changes that are poised to alter Ca^2+^ permeation and polyamine regulation. For GluE1αA, the emergence of a serine served to reduce both polyamine block and Ca^2^+ permeation. Based on NQR mutation experiments of the Kainate receptor GluR6 [59], the presence of a glycine in the NQR site, as observed on *M. leidyi* Epsilon receptors, would similarly decrease polyamine block, while a phenylalanine, present in the *B. lanceolatum* GluE1 receptor, would retain strong polyamine block. Interestingly, mammalian AMPA and Kainate receptors also exhibit adaptations in polyamine regulation, in this case via complexing with corresponding ancillary subunits (reviewed in [11]). Although the physiological significance of variations in polyamine regulation are unclear, reduced polyamine block renders channels that are more active during bouts of excitation when the membrane voltage is depolarized. Hence, while receptors with strong rectification are constrained to depolarize the membrane from negative resting voltages, those with reduced rectification can operate at a broader range of membrane voltages. Given their non-selective nature, this would increase the input resistance of the cell membrane to possibly dampen fast electrical impulses through conduction block.

## Materials and Methods

### Identification and cloning of T. adhaerens iGluRs

*T. adhaerens* ionotropic glutamate receptor sequences were identified by BLAST [63] searching a whole animal mRNA transcriptome assembly [21] using a set of NMDA, AMPA and Epsilon protein sequences from human and the ctenophore *Mnemiopsis leidyi* as queries. Candidate *T. adhaerens* sequences were then analyzed with SmartBlast [64] and reciprocal BLAST of the NCBI non-redundant database to confirm homology to iGluRs, InterPro [65] to predict conserved domains, and TMHMM [66] to predict transmembrane helices. This identified 13 iGluR sequences, 11 of which contained complete protein coding sequences and a minimum of three predicted transmembrane helices. To verify the mRNA sequences of these identified full-length transcripts and confirm their expression *in vivo*, we used nested PCR to amplify their protein coding sequences from a whole animal cDNA library utilizing primers listed in Table S5. PCR was successful for 10 of the 11 receptors, and the corresponding DNA amplicons were cloned into the mammalian expression vector pIRES2-EGFP (Clontech) using restriction enzymes listed in Table S5, followed by sequencing and analysis of triplicate independent clones to distinguish polymorphisms from PCR errors and determine consensus protein sequences for each receptor (submitted to NCBI with accession numbers listed in Table S5).

### Phylogenetic analyses of iGluR protein sequences

We started by compiling a set of eukaryotic proteomes with a balanced sampling of 88 species from the clade Amorphea (including groups such as animals, choanoflagellates, and fungi) and 96 species from Diaphoretickes (including plants and stramenopiles). Details about the included species, sources of proteomes, and their BUSCO quality metrics [67] are provided in Table S1. To generate a species aware protein phylogeny of iGluR sequences with GeneRax [31], we first created separate species and iGluR protein phylogenies which are used as inputs for the program. For the former, single copy BUSCO genes for each species were used to build a concatenated supermatrix, by first aligning homologous gene sequences with MAFFT [68] using default parameters, then trimming these with trimAl [69] using the gappyout mode, and finally concatenation with FASconCAT [70]. The species tree was built with maximum likelihood using IQ-TREE 2 [71] with the evolutionary model LG+G4+F. Branch supports were assessed with ultrafast bootstrap 1000 replicates [72] as well as with 1000 replicates for the Shimodaira-Hasegawa-like approximate likelihood ratio test (SH-aLRT) [73]. The resulting species tree was rooted in such a way as to maintain the monophyly of the Amorphea clade and was used as a backbone for gene tree-to-species tree reconciliation (see below).

Next, we used HMMER v3.1b2 [74] to generate a hidden Markov model profile of iGluRs using protein sequences from the InterPro entry IPR001320. The HMM profiles were then used to search the noted proteomes for candidate iGluR protein sequences using an expect value (e-value) threshold of 1E-10, and these were then filtered to remove redundant sequences using CD-HIT [75] with a threshold of 0.95 (*i.e.*, 95% sequence identity), except for placozoan species for which we used a threshold of 1.00 in order to retain potential transcript variants. The remaining sequences were clustered by sequence similarity using the program CLANS [76] with the alignment scoring matrix BLOSUM62 and an e-value threshold of 1E-14. The sequences from a high confidence iGluR cluster, defined by the stringent e-value 1E-65, were extracted and combined with the sequences from the InterPro iGluR entry to create a new, more refined profile with HMMER. This additional step was carried out because InterPro sequences are primarily derived from model organisms. By incorporating sequences collected from the initial HMMER search across a broad set of organisms, we sought to develop a more inclusive profile that was more likely to identify sequences in distantly related non-model organisms. A second round of HMMER with an e-value of 1E-10 was run with these updated profiles. For each species, CD-HIT was run with an identity threshold of 95% (and 100% for placozoan species) to reduce potential transcript variants from the second HMMER search. The resulting sequences were combined with sequences from the first round and CD-HIT was used with 100% identity in all cases to get rid of duplicates from the two rounds of HMMER. The resulting sequences were again clustered using CLANS using the same parameters as before. Connected sequences from the iGluR cluster with a p-value cut-off of 1E-60 (Supplementary Figure S3 and Supplementary File 2), were extracted. An additional filtering of the dataset was carried out to remove sequences with less than two or more than six transmembrane helices predicted with the program Phobius [77]. These extracted iGluR sequences were then aligned with MAFFT using the E-INS-i algorithm, and the alignment trimmed with trimAl in using the gappyout mode. The gene tree was inferred with maximum likelihood using IQ-TREE2, with a best-fit model according to Bayesian Information Criterion of Q.pfam+I+R10. Branch supports were assessed with 1000 ultrafast bootstrap replicates as well as 1000 SH-aLRT replicates. Any potential polytomy in the tree was randomly resolved using ETE3 [78]. The resulting gene tree was used as starting tree for gene tree to species tree reconciliation using GeneRax v2.1.2 [31] set to account for duplication and loss events (UndatedDL model). The utilized iGluR sequences, in raw, aligned, and aligned trimmed format, along with IQ-TREE 2 and GeneRax output files are deposited in the Zenodo repository https://zenodo.org/doi/10.5281/zenodo.11521256.

To generate a comprehensive phylogeny exclusively of placozoan iGluRs, we manually extracted candidate protein sequences from the gene data available for four species: *T. adhaerens* [20–22], *Trichoplax* species H2 [22], *Hoilungia hongkongensis* [23], and *Cladtertia collaboinventa* [24]. This was done by first BLAST searching the various databases using the *T. adhaerens* protein sequences identified in the transcriptome as queries (described above; Table S2), followed by manual annotation as described above using SmartBlast, InterPro, and TMHMM. We excluded protein sequences with 1 or 0 predicted transmembrane helices, resulting in the identification of 15 sequences for *T. adhaerens*, (11 from the whole animal transcriptome plus 3 from the Ensembl and NCBI databases and 1 comprised of merged overlapping sequences from the transcriptome and Ensembl), 12 for *Trichoplax* species H2, 10 for *H. hongkongensis*, and 12 for *C. collaboinventa* (Table S2). The identified were aligned with MUSCLE [79], and the alignment trimmed with trimAl [69] using a gap threshold of 0.6. This trimmed alignment was then used to infer a maximum likelihood phylogenetic tree using the program IQ-TREE2 [71] with a best-fit model of Q.yeast+I+G4 identified under Bayesian Information Criterion, and 1000 ultrafast bootstrap replicates to generate node support values. Top cellular expression for the different placozoan receptors was achieved by extracting single cell RNA-Seq Umifrac values for each receptor transcript [24], and determining the average value for each cell type. The placozoan iGluR sequences, in raw, aligned, and aligned trimmed format, along with IQ-TREE 2 output files are deposited in the Zenodo repository https://zenodo.org/doi/10.5281/zenodo.11521256.

### Whole cell patch clamp electrophysiology of GluE1αA expressed in CHO-K1 cells

As noted above, the cDNA of the *T. adhaerens* GluE1αA receptor was cloned into the mammalian expression vector pIRES2-EGFP, using a 5’ nested primer that inserted a mammalian consensus Kozak sequence of GCCGCCACC<u>ATG</u> (Table S5). Several mutants/variants of this receptor where also prepared via site directed mutagenesis using standard procedures and primers listed in Table S5. For electrophysiological comparisons, we commissioned GenScript (USA) to synthesize the cDNA of the human AMPA-2 heterotetrametric receptor (NCBI accession number NP_001077088.2), which was cloned into the pIRES2-EGFP vector with a Kozak sequence flanking the start codon and restriction enzymes *Sal*I and *BamH*I at the 5’ and 3’ ends of the cDNA, respectively. Receptor cDNAs in the various pIRES2 constructs were transfected into Chinese Hamster Ovary-K1 (CHO-K1; Sigma) cells cultured in Dulbecco’s Modified Eagle Medium Nutrient Mixture F-12 (DMEM/F12) supplemented with 10% Fetal Bovine Serum (FBS) and 1% penicillin-streptomycin at 37°C. Transfections were done with 2 μg of plasmid vector using the transfecting reagent PolyJet^TM^ (SignaGen Laboratories, USA) according to the manufacturer’s instructions and plated onto glass coverslips in 35 mm dishes the next day. Transfected cells were incubated at 28°C for 2 to 4 days, then coverslips were transferred to a new 60 mm dish with 5 mL of external recording solution. For the dose response, recovery from desensitization, and pharmacology experiments, the external recording solution contained: 145 mM NaCl, 4 mM KCl, 2 mM CaCl_2_, 1 mM MgCl_2_, 10 mM HEPES (pH 7.4 with NaOH; 313 mOsm/Kg). The internal solution contained: 145 mM CsCl, 2.5 mM NaCl, 1 mM EGTA, 4 mM Mg-ATP, 10 mM HEPES (pH 7.2 with CsOH; osmolarity adjusted to 304 mOsm/Kg with D-glucose). For the ion selectivity experiments, the external solution contained: 150 mM XCl ion (where X = Na, K, Li, or Cs), 10 mM tetraethylammonium chloride (TEA-Cl), and 10 mM HEPES (pH 7.4 with XOH), and the internal solution contained: 150 mM NaCl, 10 mM EGTA, 10 mM TEA-Cl, and 10 mM HEPES, 4 Mg-ATP (pH 7.2 with NaOH). For calcium selectivity experiments, the internal solution consisted of 100 mM XCl, 10 mM EGTA, 10 mM TEA-Cl, 10 mM HEPES, and 4 mM Mg-ATP (pH 7.2 with XOH; osmolarity adjusted to 326 mOsm/Kg). The external solution contained: 4 mM CaCl_2_, 155 mM TEA-Cl, 10 mM HEPES (pH 7.4 with TEA-OH; osmolarity adjusted to 326mOsm/Kg). Agonists and pharmacological compounds used in this study were dissolved in corresponding external solutions. Whole-cell macroscopic currents were recorded using an Axopatch 200B amplifier coupled to a Digidata 1550A digitizer, using the pClamp 11 software (Molecular Devices, USA). Patch pipettes were pulled using a P-1000 micropipette puller (Sutter, USA), from thick-walled borosilicate tubing (1.5 and 0.86 outer and inner diameter, respectively), to a resistance of between 1 and 6 megaohms. EC_50_ and IC_50_ values for dose-response curves were determined by fitting monophasic dose-response curves over the data using the software package Origin 2017 (OriginLab) and the equation below.

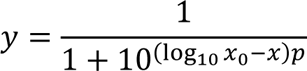

Where y is normalized current, x is the log_10_(ligand concentration), x_0_ is the EC_50_ and p is the Hill slope. The EC_50_ and IC_50_ values reported for m2 variant of GluE1αA was fitted with a biphasic dose response curve with one stimulatory and one inhibitory phase using the automated curve fitting software Dr-fit [80].

Time constant for the recovery from desensitization was calculated by fitting mono-exponential curves over the data using Origin with the following equation:

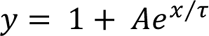

Where y is the peak current normalized to pre-pulse, A is the amplitude, x is the interval between pre-pulse and test pulse and τ is the time constant.

The relative permeability pX^+^/pNa^+^ determined for TGluE1αA, where X^+^ is a monovalent cation (Na^+^, Li^+^, Cs^+^, or K^+^), was determined using the biionic equation [81]:

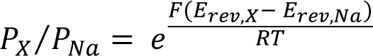

Where F is the Faraday’s constant (F = 96485.33 C/mol), R is the gas constant (R = 8.314463 J/K· mol), T is the ambient temperature (T = 294 K), and E_rev_ is the reversal potential.

The relative permeability ratio pCa^2+^/pX^+^ was determined using the following monovalent/divalent biionic equation [82]:

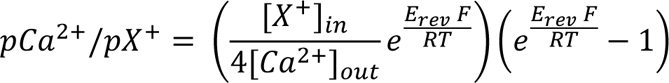

### Structural modelling and docking

The structure of the wildtype GluE1αA receptor, as well as various mutants in the ligand binding and pore domains, were predicted with AlphaFold2 [37]. For spermine docking experiments, predicted pore domains were opened in silico as described in the results. Additionally, the staring conformation of the ring of engineered asparagines in models of the S_642_N variant of GluE1αA was biased by imposed inter-subunit hydrogen bonds [83], where each asparagine donated two bonds. This bias ensured orientation of the sidechain oxygen atoms towards the pore axis. It should be noted that this orientation was imposed only once to the apo-channel model, but in the spermine-channel model, sidechain asparagines were free to move during energy minimizations. Energy optimizations and ligand docking were performed with the program ZMM (www.zmmsoft.ca) as described in a recent study [84]. Briefly, energy was calculated with the AMBER force field [85, 86], distance-and environment-dependent dielectric function [87] and implicit solvent [88]. The energy was optimized by Monte Carlo (MC) energy minimizations [89]. During MC minimizations, we used “pin” constraints to ensure the similarity of the backbone conformation in the model and the starting structure. A pin is a flat-bottom parabolic energy function that imposes energy penalties if an alpha carbon atom in the model deviates by more than 1 Å from the starting position and imposes and energy penalty with the force constant of 10 kcal·mol^−1^· Å^−2^.

## Acknowledgments

We thank Denis Tikhonov for helpful discussions. Computations were performed using facilities provided by Compute Ontario (https://www.computeontario.ca/) and the Digital Research Alliance of Canada (https://www.alliancecan.ca).

## Author Contributions

Conceived the project: A. Senatore and A. Singh; Designed and planned experiments: A. Singh, A. Senatore, B.S. Zhorov, L.A. Yanez-Guerra, and A. Aleotti; Molecular Biology: A. Singh, C.D. Yanartas, and A. Senatore; Electrophysiology experiments: A. Singh; Pharmacology experiments: A. Singh and Y. Song; Structural modeling and docking: B.S. Zhorov, A. Senatore; Phylogenetics: L.A. Yanez-Guerra, A. Aleotti, and A. Senatore; Manuscript writing: A Senatore and A. Singh; Revisions of manuscript: A Senatore, A. Singh, L.A. Yanez-Guerra, A. Aleotti, and B.S. Zhorov.

---

**Fig. S1.**
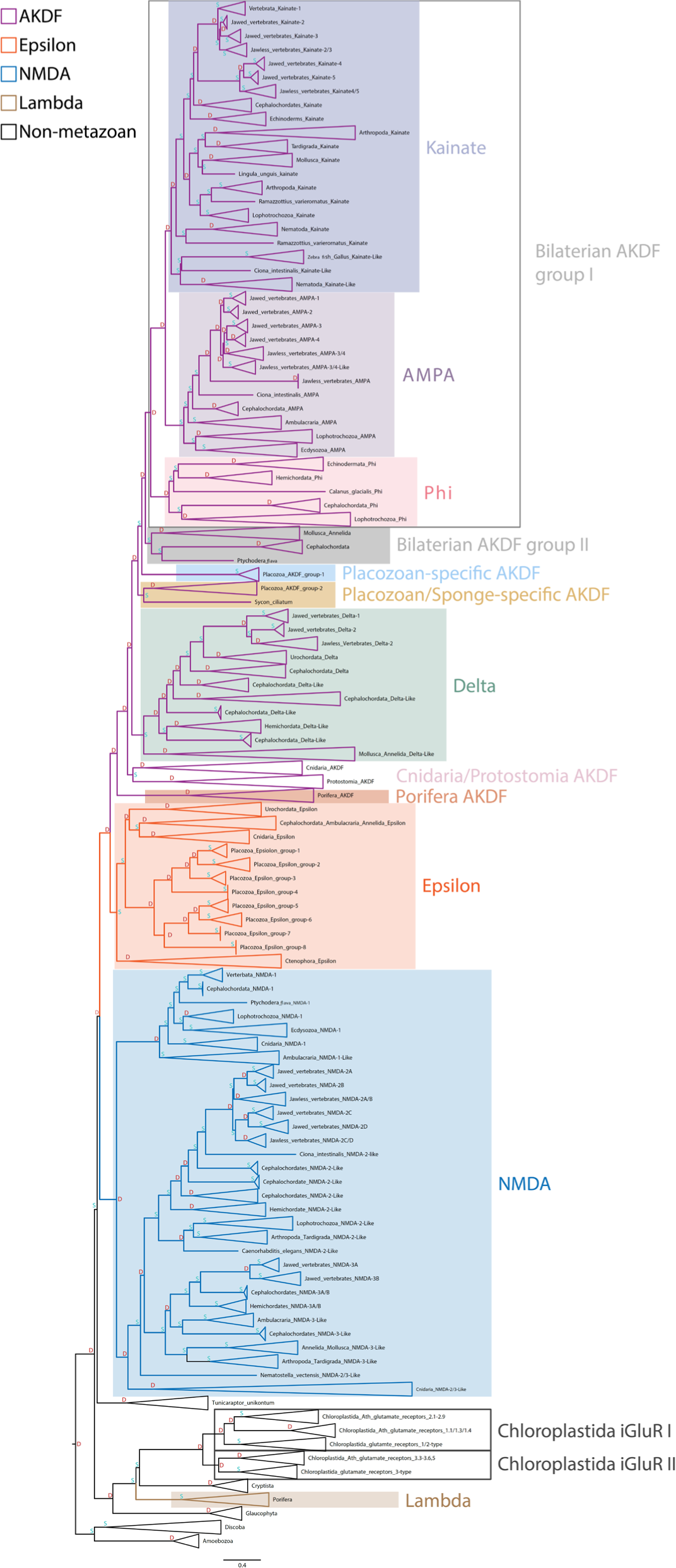
Expanded GeneRax tree of iGluR protein sequences. The letters D (red) and S (cyan) denote predicted duplication and speciation and events, respectively, predicted by the GeneRax software [31].

**Fig. S2.**
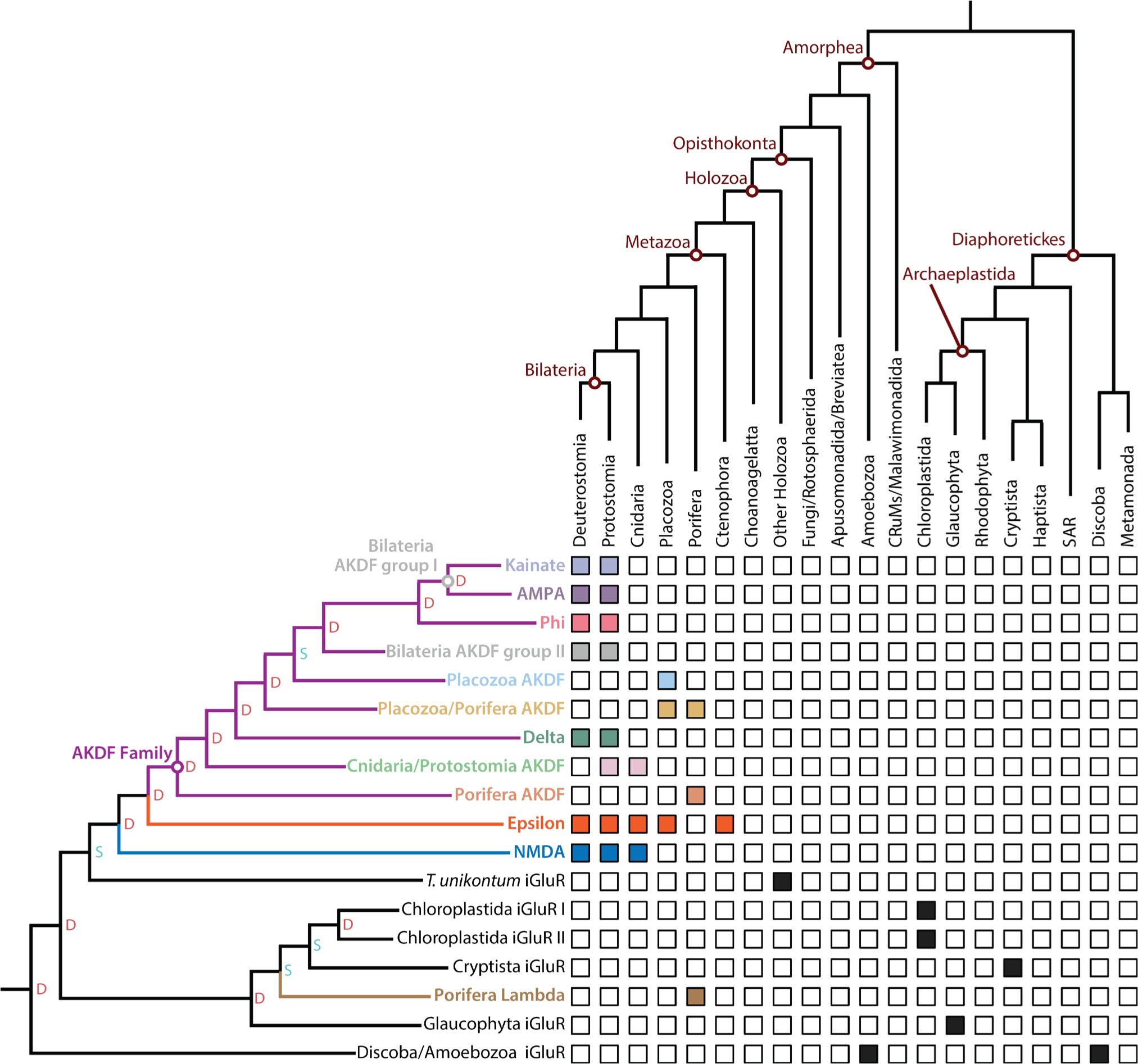
Illustration denoting presence/absence of phylogenetically distinct iGluR types in metazoans and select non-metazoan eukaryotes. The letters D (red) and S (cyan) denote predicted duplication and speciation and events, respectively, predicted by the GeneRax software [31].

**Fig. S3.**
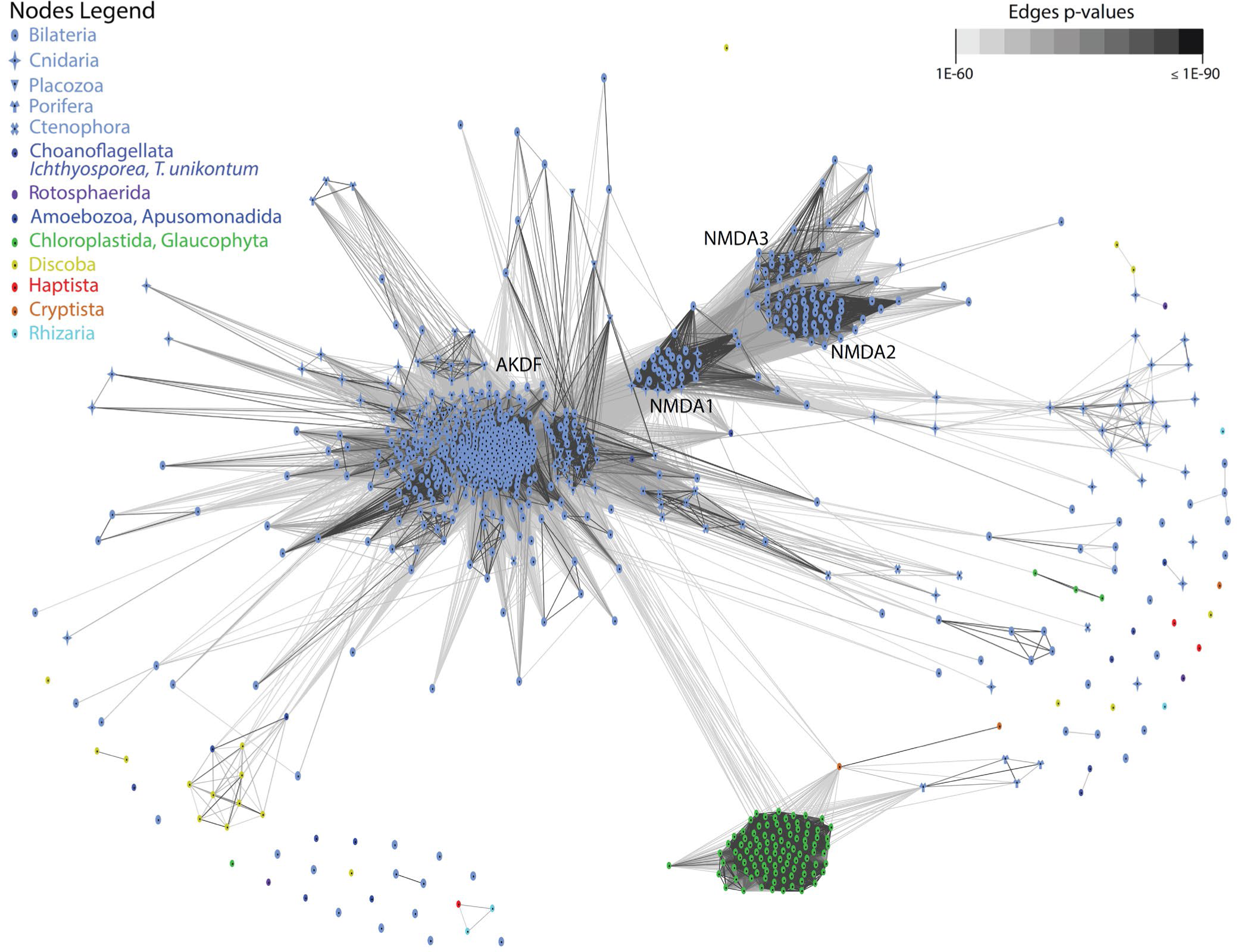
Clustering of iGluR sequences with CLANS. Dots represent sequences, colour-coded by species group. Connecting lines represent BLAST scores. Clustering is shown at p-value 1.00E-60, which was the threshold utilized to delineate the iGluR family. A large cluster of animal iGluRs (light blue) is connected to a Chloroplastida iGluR cluster (green). Four sponge sequences connect to the latter but not to the former. A cluster of Amoebozoa/Discoba sequences connect to the animal cluster.

**Fig. S4.**
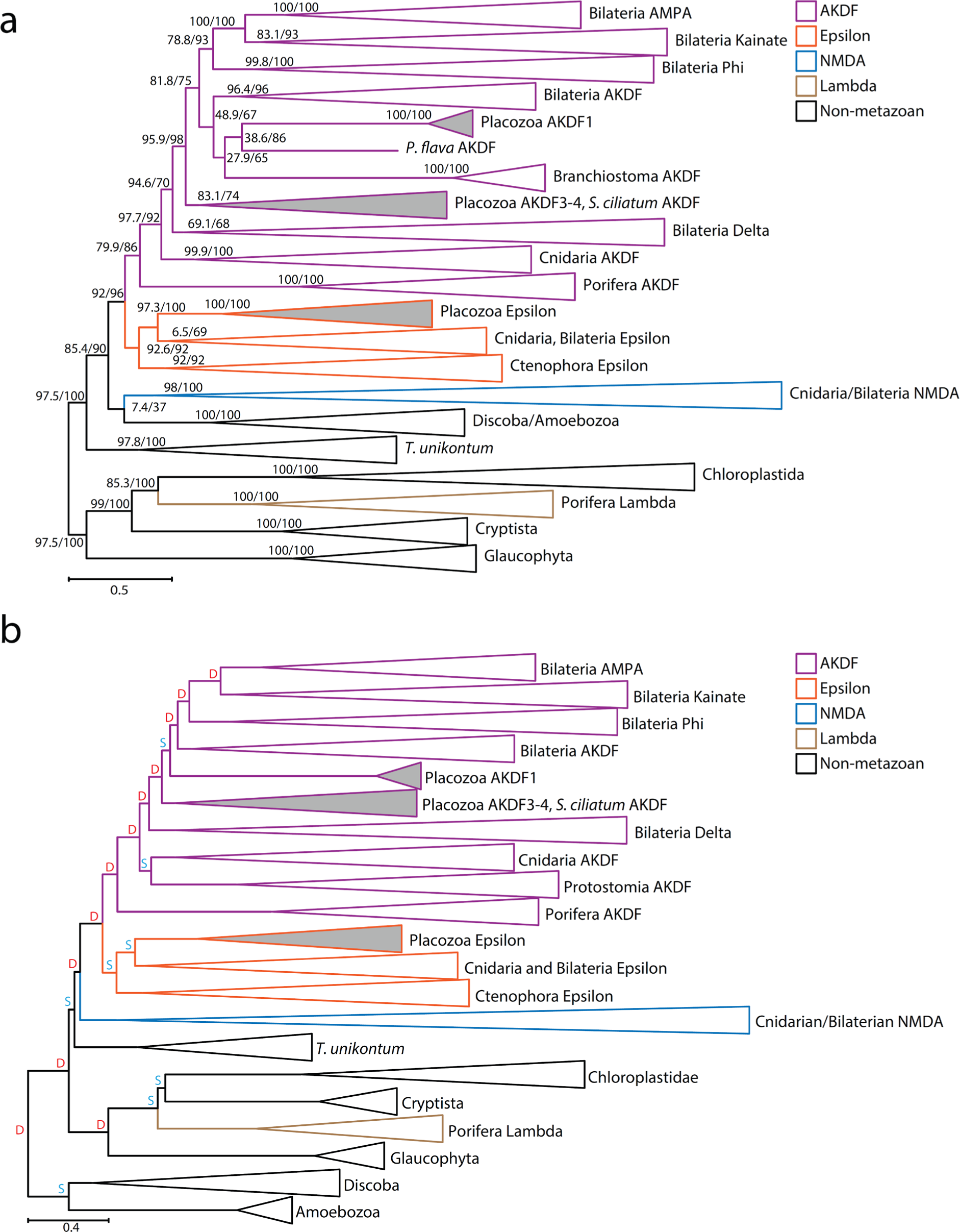
Comparison of the GeneRax species aware tree with maximum likelihood iGluR protein phylogeny. **a** Maximum likelihood iGluR tree used as starting tree for gene tree to species tree reconciliation. The tree was inferred using IQ-TREE2 [71] with the evolutionary model Q.pfam+I+R10. Branch supports are percentages for 1000 replicates for the Shimodaira–Hasegawa-like approximate likelihood ratio test (SH-aLRT) and 1000 replicates for Ultrafast Bootstrap (UFB), respectively. **b** Copy of species aware phylogenetic tree shown in Figure 1a. The letters D (red) and S (cyan) denote predicted duplication and speciation and events, respectively, predicted by the GeneRax software [31].

**Fig. S5:**
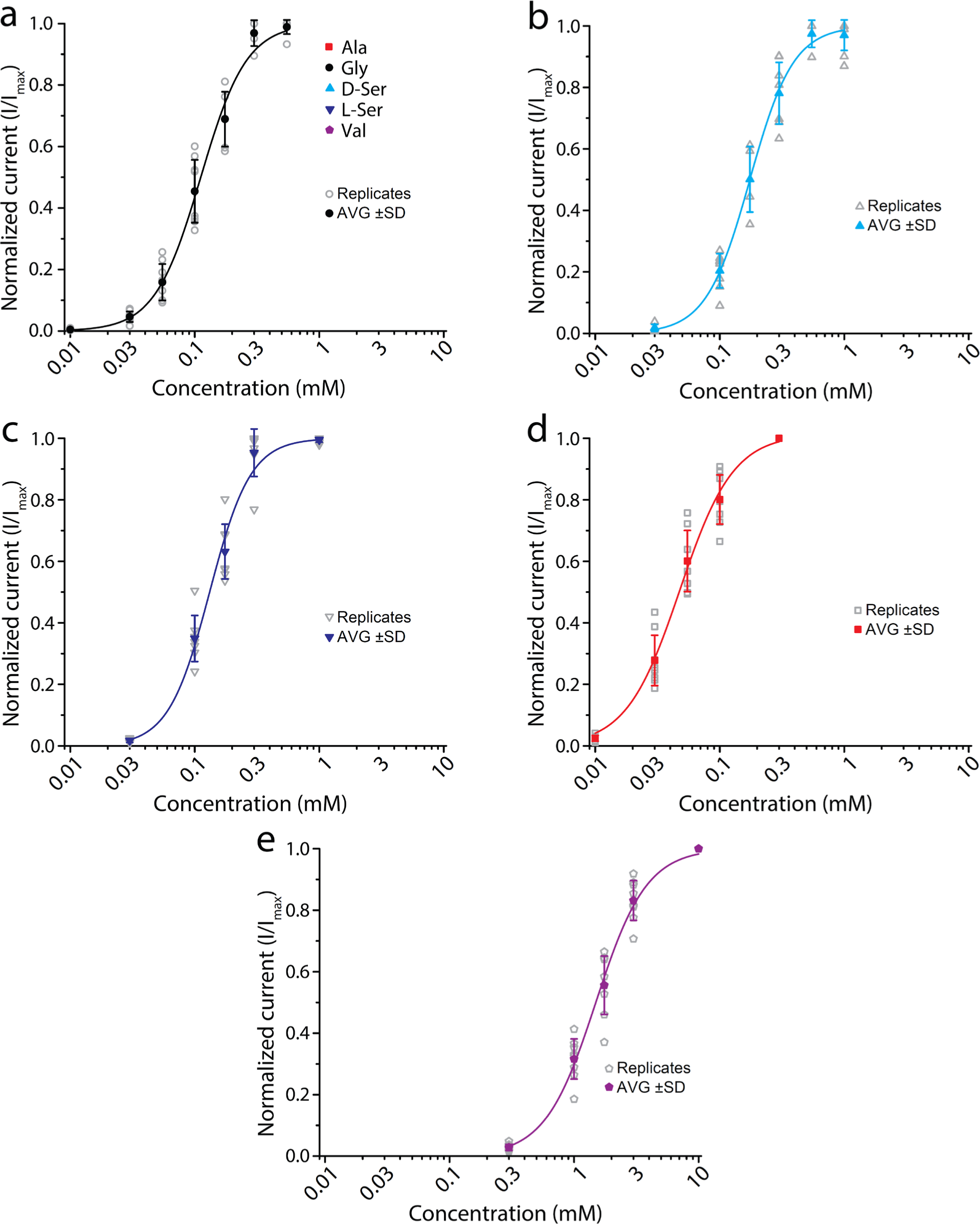
Separate dose response curves for activating ligands of GluE1αA. **a** Glycine dose response curve (n=8). **b** D-serine dose response curve (n=7). **c** L-serine dose response curve (n=7). **d** Alanine dose response curve (n=7). **e** Valine dose response curve (n=7).

**Fig. S6:**
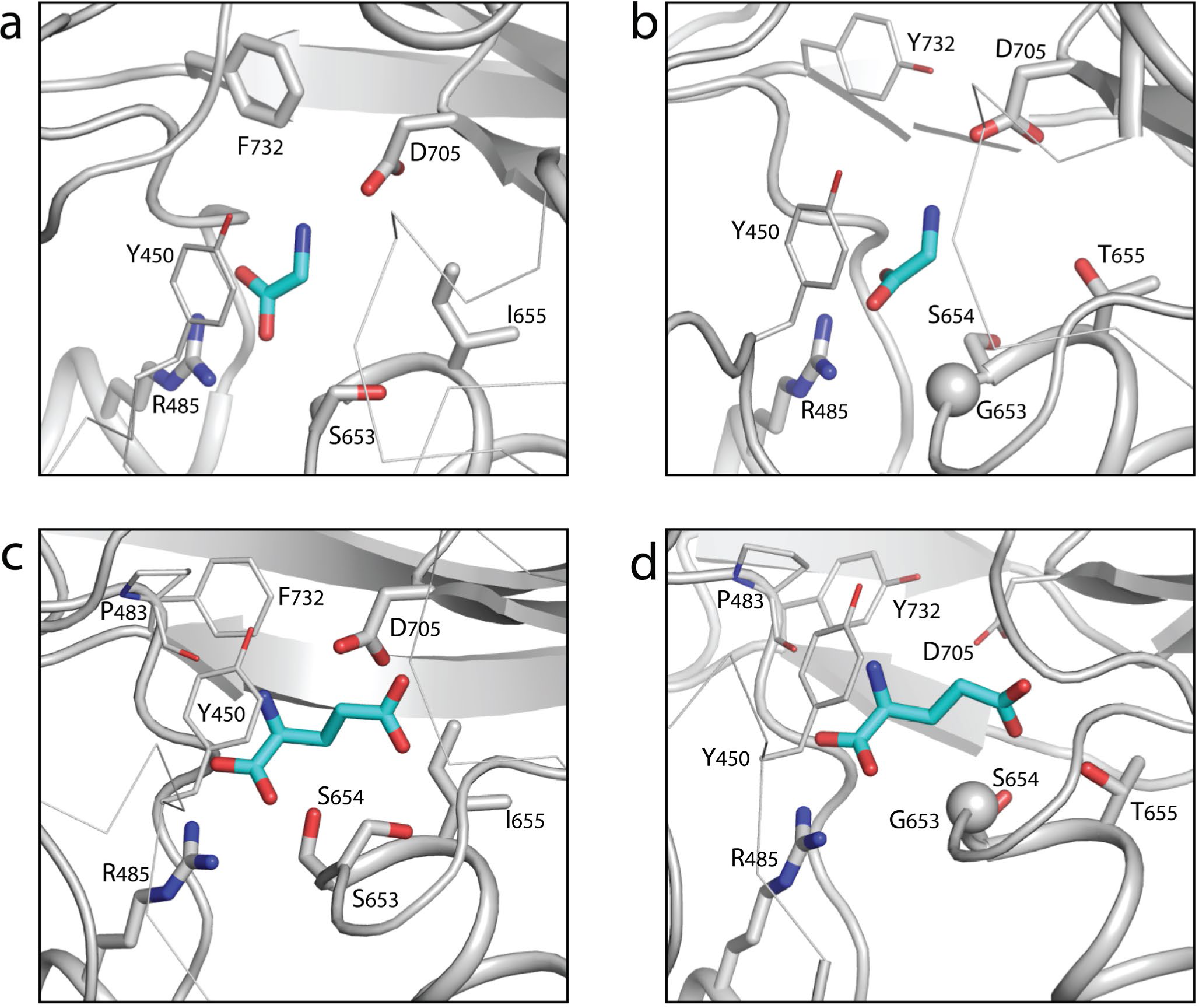
Alternate depiction of docked glycine and glutamate ligands in the ligand binding pocket of wildtype and m3 mutant GluE1αA. **a** Homology modeling and docking of glycine in the putative ligand binding pocket of the wiltype GluE1αA receptor. **b** Homology modeling and docking of glycine in triple mutant (m3) variant of GluE1αA. **c** Homology modeling and docking of glutamate in the wt GluE1αA receptor. **d** Homology modeling and docking of glutamate in triple mutant (m3) variant of GluE1αA.

**Fig. S7:**
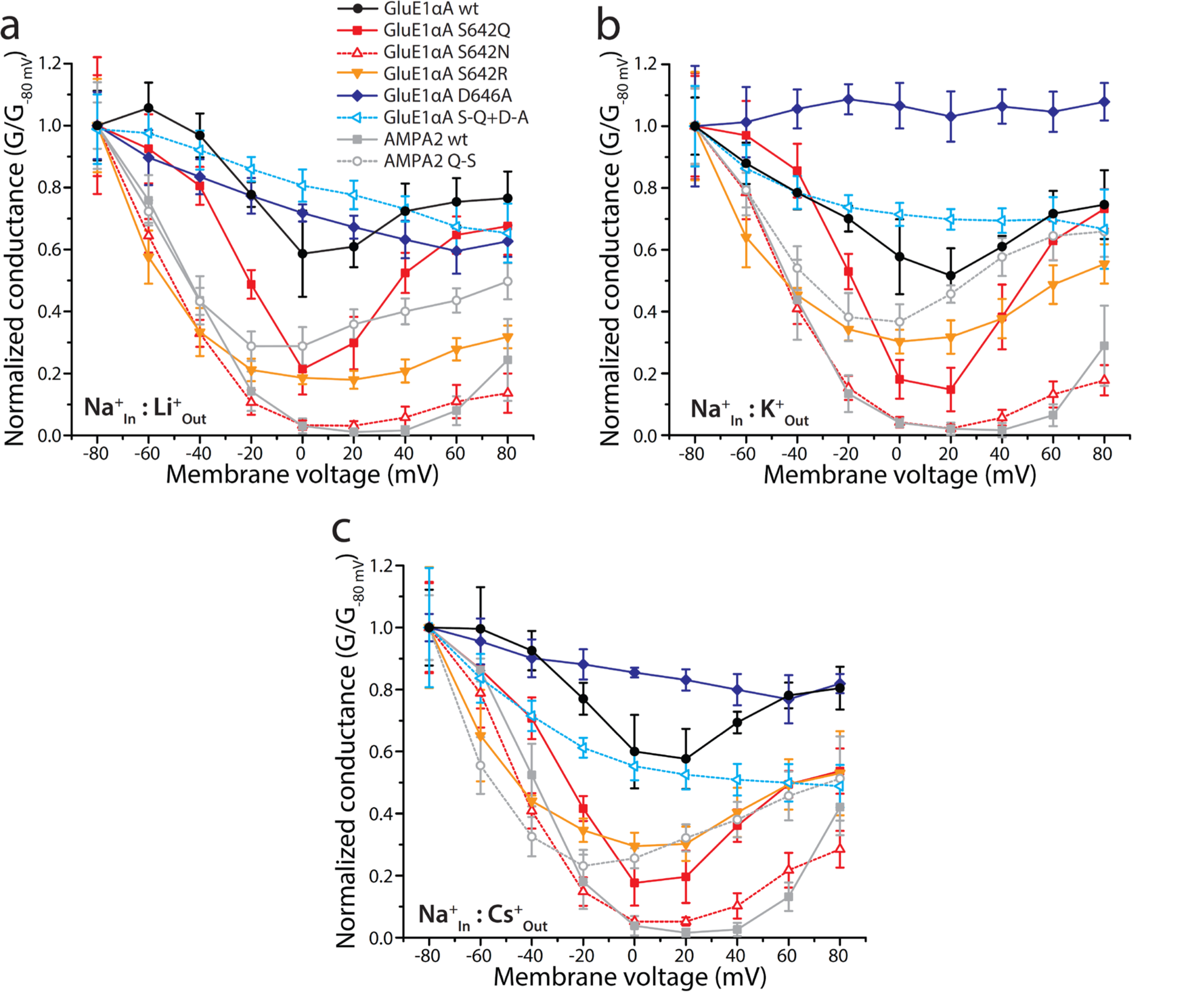
Average normalized conductance vs. membrane voltage (G-V) plots for the remaining bi-ionic conditions presented in. Fig. 5**. a** Plot of average normalized G-V data recorded using equimolar [Li^+^]_out_ and [Na^+^]_in_ for the different variants of GluE1αA and human AMPA2 receptors. **b** G-V plot for equimolar [K^+^]_out_ and [Na^+^]_in_. **c** G-V plot for equimolar [Cs^+^]_out_ and [Na^+^]_in_.

**Fig. S8:**
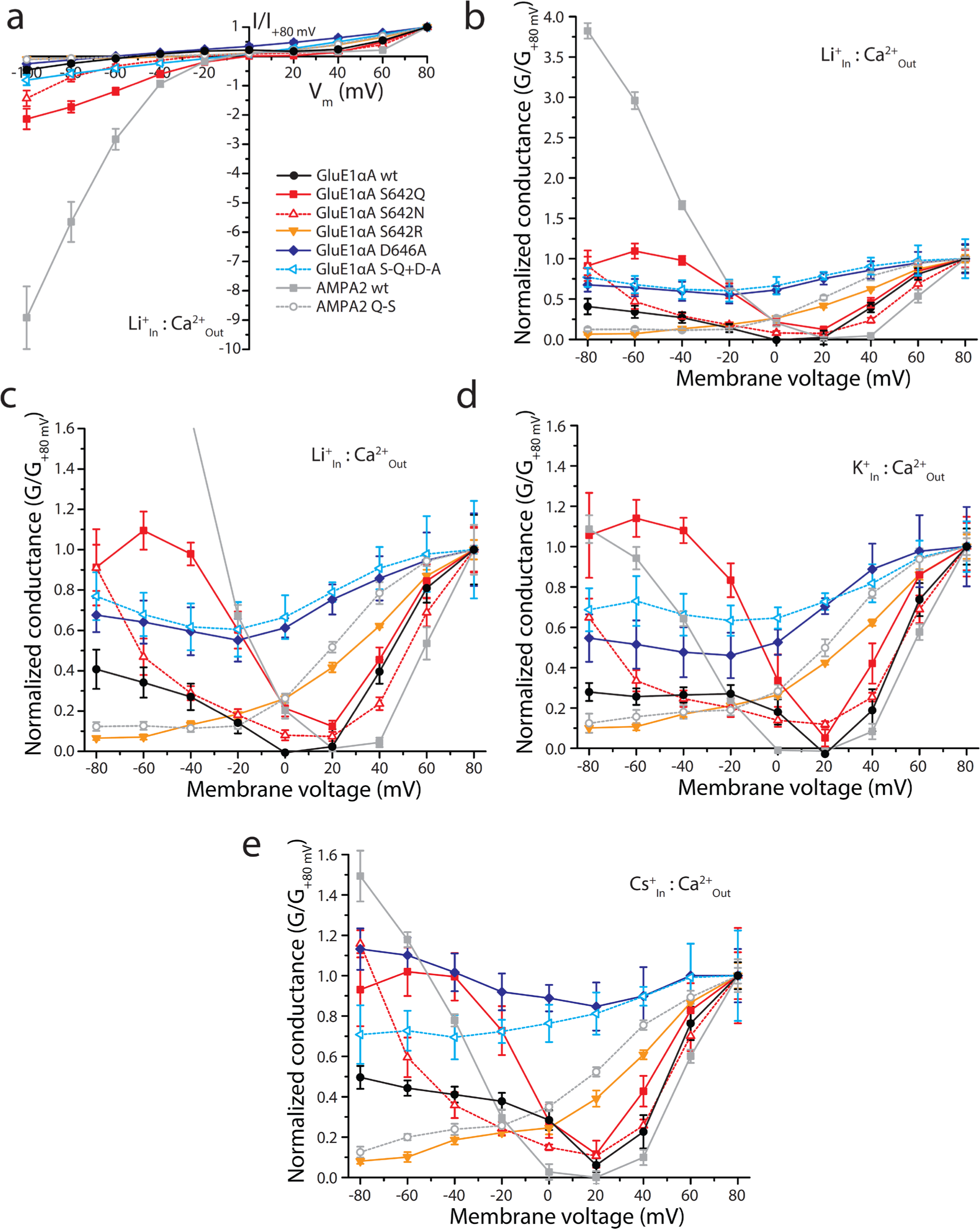
Expanded I-V plot for Li^+^ vs. Ca^2+^ and average normalized G-V plots for the remaining bi-ionic conditions presented in Fig. 6. **a** Plot of average normalized current vs. membrane voltages (I-V) under 4 mM extracellular Ca^2+^ and 100 mM intracellular Li^+^ conducted by wildtype and pore mutant variants of GluE1αA, the wildtype human AMPA2 receptor, and a mutant AMPA2 variant bearing a glutamine to serine mutation in the QRN site (n=7-8). **b** Plot of average normalized G-V data recorded using 4 mM [Ca^2+^]_out_ and 100 mM [Li^+^]_in_ for the different variants of GluE1αA and human AMPA2 receptors. **c** Scaled plot of average normalized G-V data for the plot in panel b. **d** G-V plot for 4 mM [Ca^2+^]_out_ and 100 mM [K^+^]_in_. **e** G-V plot for 4 mM [Ca^2+^]_out_ and 100 mM [Cs^+^]_in_.

**Fig. S9.**
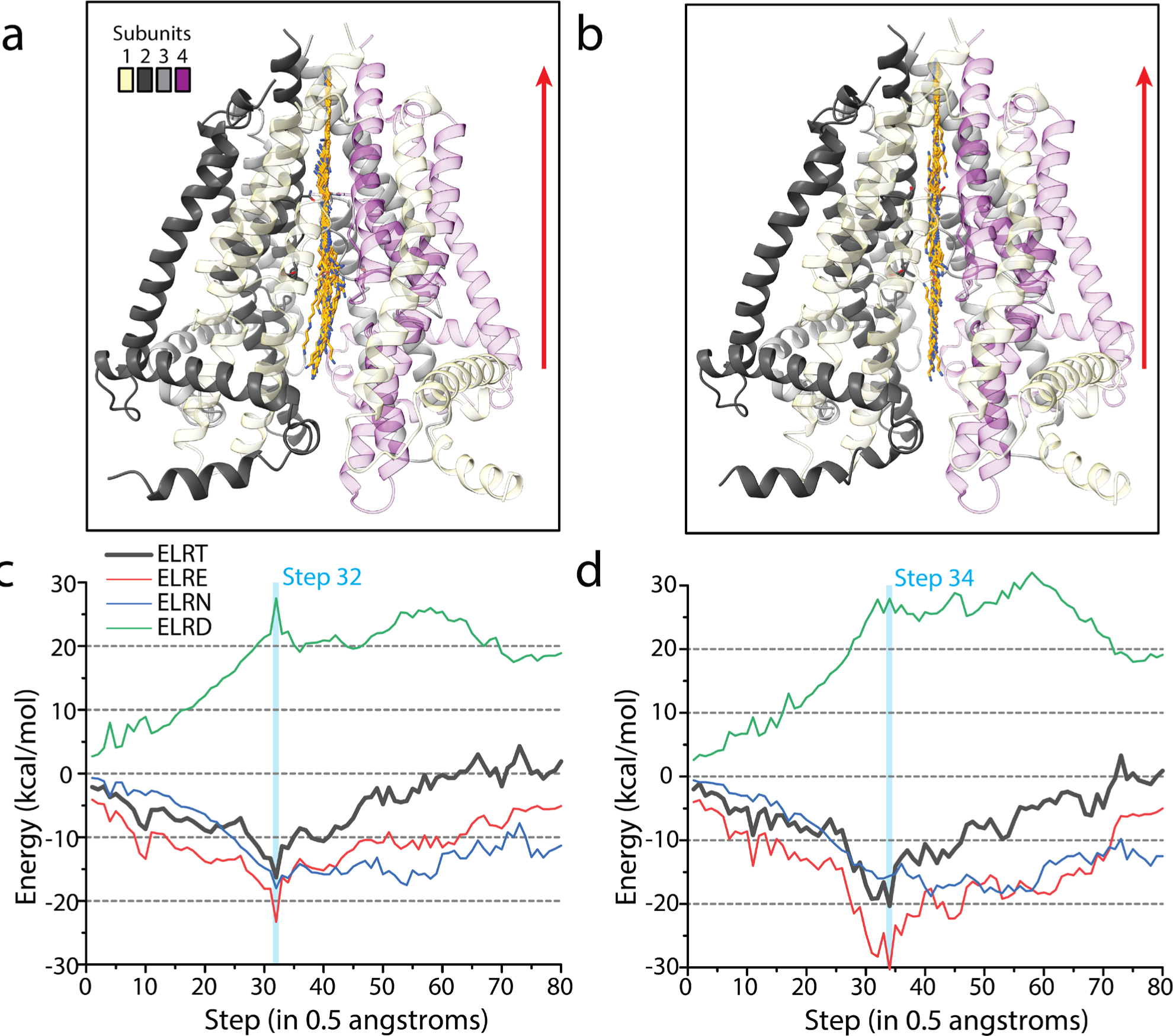
Spermine in the pore of GluE1αA. **a** Superposition of 80 MC-minimized structures of spermine within the wildtype receptor (only a single MC-minimized structure of the receptor is shown for clarity). **b** Similar superposition of 80 MC-minimized spermine structures in the N_642_N variant. For panels a and b, side chains of NQR and +4 residues are shown as sticks. Spermine is shown in orange carbons and colored heteroatoms. The front subunits 1 and 4 of the receptors are semi-transparent for clarity. The red arrows show the direction in which spermine was pulled. **c** Plot of Monte Carlo-minimized energy as spermine was pulled through the open wildtype GluE1αA pore, from the cytoplasmic to the extracellular side, in 0.5 Å steps. **d** Plot of Monte Carlo-minimized energy as spermine was pulled through the open the S_642_N variant pore. For panels c to d, ELRT: Energy Ligand Receptor - Total; ELRE: Energy Ligand Receptor - Electrostatic; ELRN: Energy Ligand Receptor - Non-bonded; ELRD: Energy Ligand Receptor - Desolvation.

