## Supplementary File 1 for "Evolution of iGluR ligand specificity, polyamine regulation, and ion selectivity inferred from a placozoan Epsilon receptor"

```
Supplementary File 1: Global alignment of T. adhaerens GluE1aA and mature rat GluA2 (UniProt ID number P19491)


  Residues listed in Table S3 are back-colored light purple.

GE1aA  1    MVKSNFIYYLLLFQAAGLIFAVTANVETTYRLGILLPRGYTKIERAIQLAIDMINENEIA  60
             V SN I                       ++G L PRG  +   A ++ +   + +E
rGA2   1    -VSSNSI-----------------------QIGGLFPRGADQEYSAFRVGMVQFSTSEF-  35

GE1aA  61   DIKLNHSRLSTTFHTADLYSPYDNFHKACNAMKEEIVAIIGPLVSGATIGAQFACSTLNM  120
                   RL+      ++ + +   +  C+     + AI G     +       C TL++
rGA2   36   -------RLTPHIDNLEVANSFAVTNAFCSQFSRGVYAIFGFYDKKSVNTITSFCGTLHV  88

GE1aA  121  PHIAP-FATDANLANNPSYTYLLRMLPVSTIESLAIASFIEYYGWTKVAILASNTDFGVS  179
              I P F TD       ++ ++++M P   ++  A+ S IEYY W K A L  ++D G+S
rGA2   89   SFITPSFPTDG------THPFVIQMRP--DLKG-ALLSLIEYYQWDKFAYLY-DSDRGLS  138

GE1aA  180  VLSKFREIASRKSWRILAFELFKIDQAGNLLDIETNLQSIKKSGARIVIINCLTSEALKI  239
             L    + A+ K W++ A  +  I+         +  Q ++    R VI++C   +   I
rGA2   139  TLQAVLDSAAEKKWQVTAINVGNINNDKKDETYRSLFQDLELKKERRVILDCERDKVNDI  198

GE1aA  240  FERAREMGMMDSGWCWIAVDAIAAETSQLP-----PNLNGLIGLAYHNNR-GKLYQNVSA  293
             ++   +G    G+ +I  +    +   L       N++G   + Y ++   K  +  S
rGA2   199  VDQVITIGKHVKGYHYIIANLGFTDGDLLKIQFGGANVSGFQIVDYDDSLVSKFIERWST  258

GE1aA  294  ----RYYLRYNETIEPLYLHYFDSVLAVAYGIQAMVREG-KLPKAAKVICDKSNPK-PWK  347
                 Y   +  TI+      +D+V  +    + + ++  ++ +        +NP  PW
rGA2   259  LEEKEYPGAHTATIKYTSALTYDAVQVMTEAFRNLRKQRIEISRRGNAGDCLANPAVPWG  318

GE1aA  348  QGHQMFHFIKKASGPSTTSSINFNANGGPSNVGYDIINLHGQSWAKVGSWTNHRNLILDR  407
            QG ++   +K+      + +I F+ NG   N   +I+ L      K+G W+    +++
rGA2   319  QGVEIERALKQVQVEGLSGNIKFDQNGKRINYTINIMELKTNGPRKIGYWSEVDKMVVTL  378

GE1aA  408  NRIYFLNGVNHILDSGSFIGG---KTLTVTTILDAPFTMADRDYAIT--GRKYRGYIIDL  462
                        L SG+   G   KT+ VTTIL++P+ M  +++ +     +Y GY +DL
rGA2   379  TE----------LPSGNDTSGLENKTVVVTTILESPYVMMKKNHEMLEGNERYEGYCVDL  428

GE1aA  463  LDEMSKNLNFTYKIRIVADGQYGSQYTDKNGALKWTGVIGEVIDGIADMAAAPLSITPER  522
              E++K+  F YK+ IV DG+YG++  D      W G++GE++ G AD+A APL+IT  R
rGA2   429  AAEIAKHCGFKYKLTIVGDGKYGARDADTK---IWNGMVGELVYGKADIAIAPLTITLVR  485

GE1aA  523  QQALDFTMPFMNQGLTVLTLTKKNEANSLFQAFLPLKIEVWIGILISLIVVAIATTCMNR  582
            ++ +DF+ PFM+ G++++    +     +F    PL  E+W+ I+ + I V++    ++R
rGA2   486  EEVIDFSKPFMSLGISIMIKKPQKSKPGVFSFLDPLAYEIWMCIVFAYIGVSVVLFLVSR  545

GE1aA  583  WSPFDYYGKAAEKLQMLEDRYQSESKLQHNYDYWQMEKEEALEAFSFGNTLWYTLGSFLS  642
            +SP++++       +  ED  +++S              E+   F   N+LW++LG+F+
rGA2   546  FSPYEWH------TEEFEDGRETQSS-------------ESTNEFGIFNSLWFSLGAFMQ  586

GE1aA  643  QGADRTPRSISARLITAIWWLSSVIIIATYTANLTAFLTVSSLQTTFNSLHDLAKSSGMG  702
            QG D +PRS+S R++  +WW  ++III++YTANL AFLTV  + +   S  DL+K + +
rGA2   587  QGCDISPRSLSGRIVGGVWWFFTLIIISSYTANLAAFLTVERMVSPIESAEDLSKQTEIA  646

GE1aA  703  YGVLANSSIEYFFLNTAVSPYQEMKHNLKN------VRSNMEGVQRVLSSTTEHYAYIGD  756
            YG L + S + FF  + ++ + +M   +++      VR+  EGV RV  S  + YAY+ +
rGA2   647  YGTLDSGSTKEFFRRSKIAVFDKMWTYMRSAEPSVFVRTTAEGVARVRKSKGK-YAYLLE  705

GE1aA  757  AAVLKYAKSQY-CNLTTVG-SFKEDSFGLALPKESLYWKEVSIQILRFREEGFLETLQTK  814
            + + +Y + +  C+   VG +     +G+A PK S     V++ +L+  E+G L+ L+ K
rGA2   706  STMNEYIEQRKPCDTMKVGGNLDSKGYGIATPKGSSLGNAVNLAVLKLNEQGLLDKLKNK  765

GE1aA  815  WF--EGKCKA----VSESTSHDPVKFESLIGVFYILLATVGFSFAILITEWIVAAIKDTR  868
            W+  +G+C +      E TS   +   ++ GVFYIL+  +G +  + + E+   +  + +
rGA2   766  WWYDKGECGSGGGDSKEKTS--ALSLSNVAGVFYILVGGLGLAMLVALIEFCYKSRAEAK  823

GE1aA  869  RHYMKISFAGAIKKRFLYIINDSFSATFCKDKTDRCNTGELSVTKTKRTSTLI  921
            R  MK++       +    IN S S       T +       +   K     I
rGA2   824  R--MKVA-------KNPQNINPSSSQNSQNFATYKEGYNVYGIESVK-----I  862
```
